# Unravelling the genetic architecture of persistence in production, quality, and efficiency traits in laying hens at late production stages

**DOI:** 10.1101/2025.02.26.640268

**Authors:** Quentin Berger, Nicolas Bedere, Sandrine Lagarrigue, Thierry Burlot, Pascale Le-Roy, Thierry Tribout, Tatiana Zerjal

**Affiliations:** Université Paris-Saclay, INRAE, AgroParisTech, GABI, 78350, Jouy-en-Josas, France; PEGASE, INRAE, Institut Agro, 35590, Saint-Gilles, France; NOVOGEN, 5 rue des Compagnons 22960 Plédran, France

**Keywords:** Persistence, genetic architecture, laying hens, egg production, egg quality, efficiency

## Abstract

**Background:** The laying hen industry aims to extend production for the economic and environmental benefits it offers, while at the same time facing the challenges of declining egg production and quality in aging hens. To explore trait persistence, we studied 998 Rhode Island Red purebred hens from the Novogen nucleus. We recorded daily egg production from 70 to 92 weeks of age and measured individual feed intake twice a week for three weeks, starting at 70, 80, and 90 weeks, as well as body weight at the start and at the end of each feed intake recording period. Random regression models were used to study trait trajectories over time, and PCA and hierarchical clustering were applied to identify groups of hens based on estimated breeding values for the intercept and slope of traits trajectory.

**Results:** Results showed different aging trajectories among traits. Daily body weight variation, Feed conversion ratio, Haugh unit and yolk percentage showed persistence (i.e., stability) over the measured period. On the contrary, daily feed intake, residual feed intake, laying rate, egg mass, eggshell breaking strength and stiffness decreased over time, while body weight, mean egg weight and eggshell colour increased. To assess the feasibility of selecting for trait persistence, we estimated the genetic variance of the slope and its correlation with the intercept. We found that, for egg weight and eggshell colour, genetic variance of the slope was negligible, indicating that selection for persistence on these traits requires other means. On the contrary, the slope for other traits such as laying rate and residual feed intake showed significant additive genetic variance. Strong genetic correlations between trait estimates at different ages were also observed and heritabilities estimates were low to high depending of the traits and period.

**Conclusion:** The study explores hens’ trait persistence from 70 to 92 weeks, suggesting potential for improved egg production persistence. Challenges arise from low genetic variances impacting the efficiency of the potential selection on persistence. Clustering analysis reveals distinctive response patterns to elongation of production and underlined that selecting for enhanced persistence of different traits will necessitate compromises in breeding goals.

## Background

The egg production sector is an important player in food security as eggs represent one of the most affordable animal protein sources, worldwide. The projected development of global egg production over the next 10 years is of 1% increase per year, leading to an expected global production of 96 million tons in 2030 [1]. The layer breeding industry is looking for strategies to conciliate a constant increase in egg demand due to a growing human population, ever larger climatic and political constraints, and a new societal environmental and animal welfare awareness.

One promising strategy implemented to maintain profitability while remaining socially responsible and environmentally friendly is the extension of the laying period beyond 70 weeks of age. The practice of extending the laying period of commercial hens can offer numerous benefits. From an economic perspective, keeping hens beyond 70 weeks of age can yield significant gains by increasing the number of saleable eggs per hen and cushioning the non-productive period. With a longer laying cycle the unproductive periods are reduced, as well as the costs related to bird replacement, while higher revenue for egg producers due to increased egg production are expected. Additionally, keeping hens for a longer period can reduce the frequency of hen replacements, depopulation, and cleaning of the poultry unit, which can translate to lower costs and fewer resources needed in the long term [2]. Ethically, extending the laying period reduces the size of flocks and the number of hens that are sacrificed at the end of the production period. As shown in Great Britain, increasing of 25 units the number of eggs per hen during the production life will lead a reduction of the number of productive hens of 2.5 million birds per annum [3], that corresponds to a decreasing of the hen population in England of 7.6% [4]. Environmentally, reducing the number of layers flocks helps decrease feed requirements and associated environmental costs, such as energy and land use [5, 6]. It would also contribute to reduce the eutrophication impact as it was estimated that nitrogen excretion decreases by around 1 g per dozen eggs for 10 weeks production extension [3].

Nonetheless, extending the production period is not without consequences, as it impacts both animal productivity and egg quality. Several studies have shown that egg production declines beyond 60 weeks of age at a rate of approximately 7 to 10 % every 10 additional weeks of laying cycle [2, 7, 8]. Therefore, at 100 weeks of age, laying rates of 65-70% are quite common in commercial laying hens [9, 10]. Another aspect to consider when extending the laying period is that egg weight increases with bird age whilst shell quality tends to deteriorate. This is evidenced by more fragile egg shell, a reduction in the Haugh unit (HU) [2, 11], and an increase in egg deformities [12]. The mean decrease in quality is also associated with an increase in the variability of egg quality [3], with nearly 25% of eggs being unmarketable [6]. Finally, body weight tends also to increase with the hen age and the gain in weight is often due to abdominal fat accumulation, which, if excessive, can negatively affect hens’ reproductive performance [13, 14]. Furthermore, as egg production is a demanding process for the overall metabolism of laying hens, extending the production period may elevate the prevalence of metabolic diseases [3].

The decades of genetic selection of commercial lines of layers to improve laying rate, has resulted in a remarkable egg production improvement [15]. Research on the genetic architecture of traits related to egg production during early production periods up to 60 weeks of age is abundant. Studies have primarily focused on genetic improvement of the number of egg laid on the production period [12, 16–19], estimating the possibility of selection directly on laying curve [20], or identifying proxies for selecting on egg quality traits [21]. On the contrary, there is a scarcity of research concerning late production periods, despite the current priority to increase egg production through breeding for increased persistence in lay at later laying cycle stages [22].

While the concept of persistence in egg-laying hens is well known and has been discussed among scientists [23], the literature on the genetic determinism of this trait remains limited [2, 20]. Previous research has mainly focused on the genetic architecture of egg production throughout the animals’ productive life, examining either the whole production period [17, 24] or specific intervals within it [18], the former being the closest to the selection objective, while the latter being theoretically the best approach [25]. As a result, improving persistence has often involved increasing the number of eggs laid during these different periods. However, this approach does not account for the trajectory of the trait over time in each individual animal, which is essential for improving trait persistence. One method for selecting animals based on traits measured sequentially is to use random regression models (RRM). These models allow for the prediction of an animal’s estimated breeding value trajectory for a given trait over time, enabling the identification of the most persistent individuals from a genetic perspective. Moreover, this approach allows selecting animals based on the complete pattern of performances rather than isolated performances at specific times, which are often treated as multiple traits [26]. Random regression models are commonly used in dairy cattle evaluations [27–33], and their application to layer performance is less common and mainly restricted to egg production [20, 25, 34, 35]. In our study we applied RRM on a variety of traits in laying chickens measured from 70 to 92 weeks of age, a production period seldom studied, to i) investigate the dynamic of genetic parameters of egg production, egg quality, and feed efficiency traits over this period of time, ii) evaluate the potential of selecting animals on genetic random regression coefficient estimates to improve traits persistence after 60 weeks of age; iii) identify specific hen types based on their traits’ persistence.

## Materials and Methods

In this study we employed random regression models to analyse the dynamics of variance components and breeding values for efficiency and production related traits over time. This approach allowed us to investigate individual reactions to prolonged production periods and identify potential strategies for selecting chickens with enhanced persistence.

### Birds and farming system

This study used 998 hens from a Rhode Island Red Line selected for egg production and quality, body weight and feather pecking, from the nucleus of the Novogen breeding company (Pledran, France). The hens were produced in two consecutive batches of 501 and 497 animals. The hens were reared on floor pens up to 17 weeks of age. After that, all hens from the first batch and 377 from the second one were transferred in a barn with collective cages of full sibs (five birds per cage), while the remaining 120 animals from the second batch were transferred in a barn with floor pens. At 55 weeks, all animals were moved to individual cages until they were slaughtered at 93 weeks of age. The collective cages had a living area containing food and water, while the floor system had nests overlaid in two rows.

The animals were fed *ad libitum* a commercial laying diet (11.47 MJ/kg metabolizable energy, 16.5% of crude protein, 3.70% of Ca) and the lighting regime was the standard 16h of light and 8h of darkness and ambient temperature in the barn was kept at 20°C constant.

To ensure precise pedigree records, we traced the relationship between animals over at least five generations. This allowed to retrace a pedigree encompassing 27390 animals, organized into sire-based families with an average of 22 offspring per sire and 5 offspring per dam. For our study, this translates to an average of 27 sires and 111 dams per batch.

### Phenotypic recording

A large range of traits were recorded between 70 and 92 weeks of age and they are summarised in Figure 1. Egg production was recorded daily from 70 weeks to 92 weeks of age. Laying rate (LR) was calculated weekly by dividing the number of eggs laid during the considered week (EggLaid) by 7 and multiplying the result by 100. Laying rates below 50% were considered as outliers and removed from the analysis. Individual feed intake (FI) was measured during three periods of three weeks each, starting at 70, 80, and 90 weeks of age (i.e., from 70 to 82 wks, from 80 to 82 wks and from 90 to 92 wks). Feed intake was recorded twice a week, every 3-4 days, by weighing the individual feeders. The weekly feed intake (WFI) and daily feed intake (DFI) were calculated as follows:

**Fig. 1:**
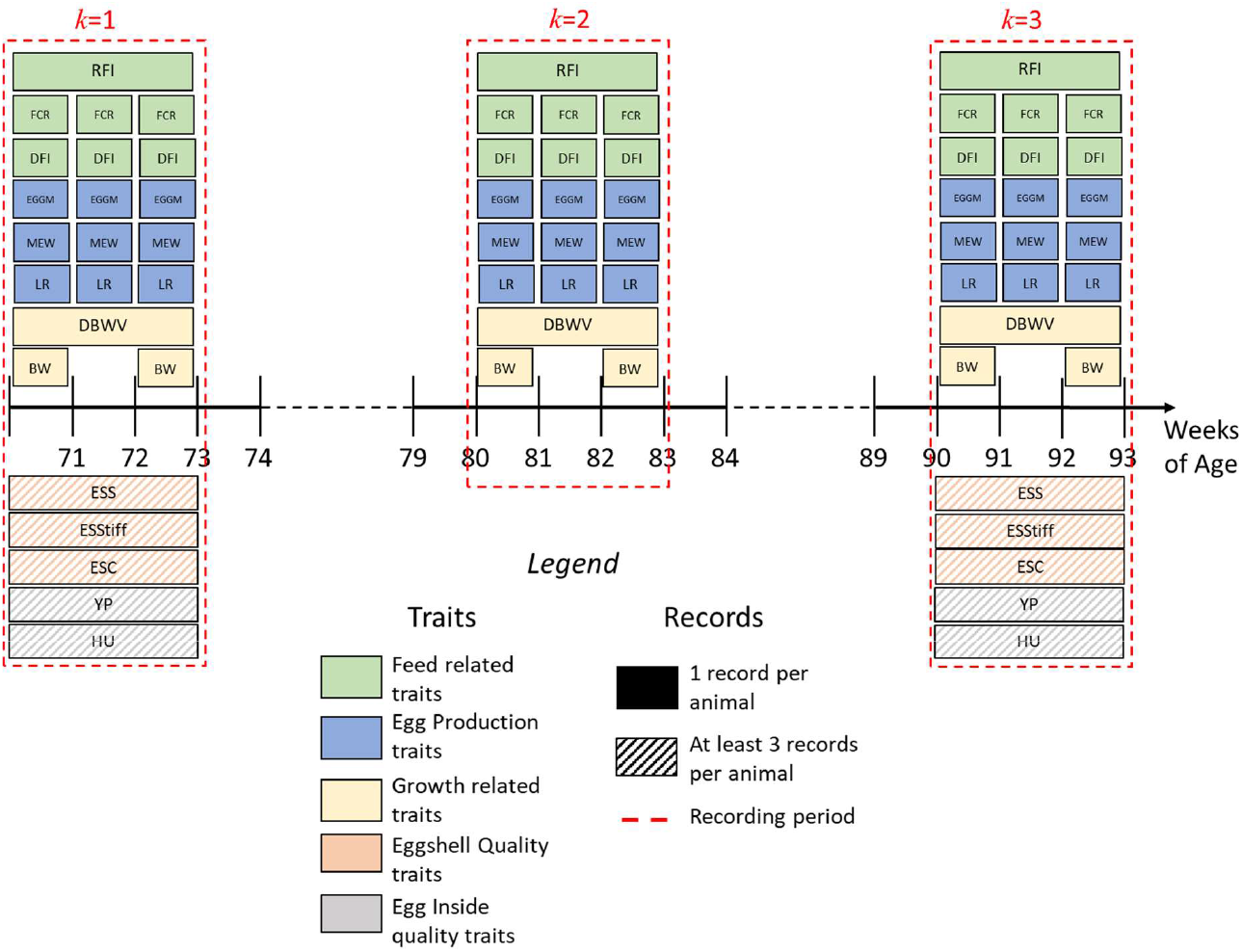
Summary of the data used in the study. Summary of the collection strategy

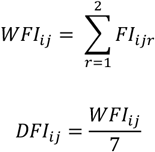

Where *FI*_*ijk*_ is the feed intake of animal *i* during the week *j* for the weekly recording records *r*.

Body weight (BW) was measured for every hen at the start and the end of each period of 3 weeks (at 70, 72, 80, 82, 90 and 92 weeks of age). The egg weight (EW) was recorded daily from 70 weeks of age to 92 wks and the mean egg weight (MEW) was determined for each week by averaging the weight of eggs laid during each recorded week.

The egg mass (EGGM) and feed conversion ratio (FCR) were calculated weekly as follows:

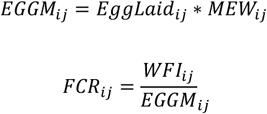

where *EggLaid*_*ij*_ is the number of eggs laid, *MEWij* the mean egg weight and *WFI*_*ij*_ the weekly feed intake for the animal *i* on the week *j*.

Daily body weight variation (DBWV) and residual feed intake (RFI) were calculated for each of the three feed recording periods. DBWV was calculated as the difference between the body weight measured at the end and at the start of the 3-week feed intake measure period, divided by 21, as described in the formula:

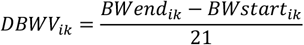

with *BWend*_*ik*_, *BWstart*_*ik*_, being the body weight for the animal *i* recorded at the end and at the start of the period *k*.

For each of three 3-week recording periods *k* and for each individual animal *i*, the Residual Feed Intake (RFI_*ik*_) was computed as the difference between the actual feed intake and the predicted feed intake for the same period 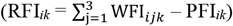.The actual feed intake for each period *k* was calculated as the sum of the weekly recorded values (WFI_*ij*_) recorded over the three weeks *j (j=*1,2,3). The predicted feed intake (PFI_*ik*_) was estimated through a multiple regression equation as described in references [36, 37]. The three independent variables recorded during each *k* period (*k*: 70-72; 2: 80-82; 3: 90-92 weeks of age) used in the equation are: mean body weight (BW_*meanik*_), DBWV_*ik*_, and egg mass (EGGM_*ik*_). The equation’s parameters underwent adjustment using the *nls* function in the R programming language [38], and the equation was fitted to the combined data of all animals over the three 3-week periods:

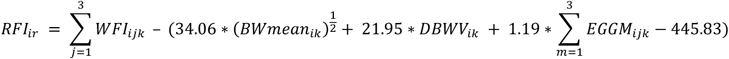

Egg quality traits were measured by the Zootests company (Ploufragan, France) on at least three individual egg records per hen per period on the first and last 3-week recording periods.

Eggshell breaking strength (ESS) and egg shell stiffness (ESStiff) were measured by compressing eggs between two flat plates moving at constant speed, using a compression machine (model Synergie 200, MTS Systems Corporation, Minnesota, United-States). ESS is the maximum force recorded before eggshell fracture. ESStiff represents deformation of the shell under a constant force of 15 Newton. Eggshell color (ESC) was obtained by recording redness (a*), yellowness (b*) and lightness (L*) with a portable chromameter (model CR400, Konica Minolta, Osaka, Japan). Yolk percentage was estimated as the ratio between yolk weight and egg weight multiplied by 100 and the Haugh unit was calculated using the formula by Haugh [39].

### Models

Weekly phenotypes over the three 3-week recording periods were analysed with random regression models, whose variance components were estimated with an average information restricted maximum likelihood (REML) method [40, 41], using the ASRemL software [42]. To do so, we modelled the trajectory of the traits with first degree polynomials, over the three recording periods, our aim being to study the rate of change of the performances rather than perfectly modelling their kinetics. The model fitted for all the traits is the following:

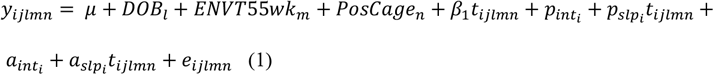

where *y*_*ijlmn*_ represents the dependent variable for the hen or egg *i* of the batch *l*, from the rearing system *m*, and cage position *n* on the week *j, μ* the intercept, *DOB*_*l*_ the fixed effect of the date of birth of animals in the batch *l, ENVT55wk*_*m*_ the fixed effect of rearing system before 55 weeks (*m* = floor or collective cage), *PosCage*_*n*_ the fixed effect of the cage location in the barn (n = 1:32, each value identifying a sequence of 16 cages), β_1_ the fixed regression coefficient of the animal’s age t (which is the environmental gradient of the reaction norm, from *t*_70w_= 0 to *t*_92w_= 22). For egg quality phenotypes an additional fixed regression coefficient, whose covariate was the time span between the day of egg lay and the day of quality measurement, was added to the model. The linear function modelling the random permanent environmental effect of the animal *i* is 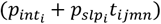,where 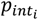 and 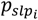 are the random intercept and random slope of the random regression on the age t, respectively, with 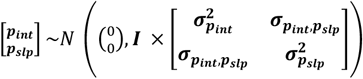
where **I** is the identity matrix, σ^2^_pint_ is the permanent environment variance for the intercept term, σ^2^_pslp_ the permanent environment variance for the slope term, and σ_pint,pslp_ the permanent environment covariance between intercept and slope term.

Similarly, 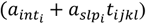 models the random additive breeding value of animal *i*, where 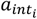 and 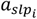 are the random intercept and random slope of the random regression on the age t, respectively, with 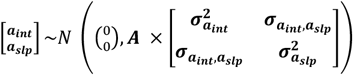,where **A** is the pedigree-based relationship matrix, σ^2^_aint_ is the additive genetic variance for the intercept term, σ^2^_aslp_ is the additive genetic variance for the slope term, and σ_aint,aslp_ the covariance between the two aforementioned effects.

Finally, *e*_*ijkl*_ is the random residual effect, with

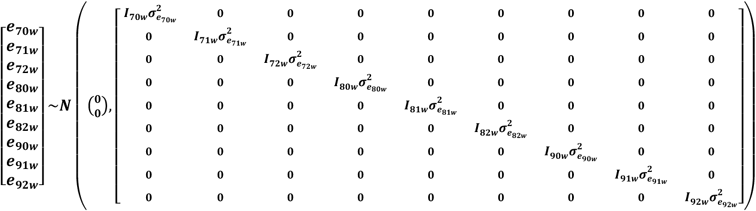

where **I**_***70w***_ to **I**_***92w***_ are the identity matrices and 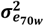 to 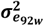 are the residual variances at the corresponding age of measure.

Significance of the genetic (co)variances has been tested by comparing our model with reduced models, by log-likelihood ratio tests, where either the genetic variances of 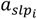 or the genetic covariance between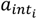 and 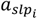 have been set to 0.

### Genetic parameters

The additive genetic and permanent environment variances for a trait at the time *t*_*1*_, (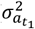 and 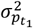,respectively), and the genetic and permanent environment covariances for a trait between two times *t*_*1*_ and *t*_*2*_, along the gradient of 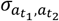 *and* 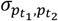,respectively), have been computed using the formulas:

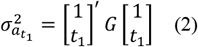

Where 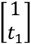 is the vector containing the covariates of the additive random regression coefficients 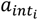 and 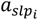 at recording time *t*_*1*_, and 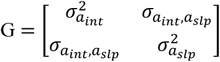 is the matrix of genetic covariance estimates between the two additive genetic random regression coefficients. Equation (2) can be explicitly stated algebraically as:

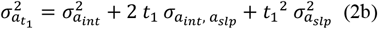

Similarly,

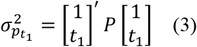

where 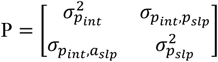 is the matrix of permanent environment covariance estimates between the two permanent environment random regression coefficients 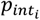 and 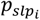.Equation (3) can be explicitly stated algebraically as:

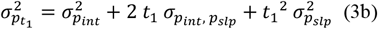

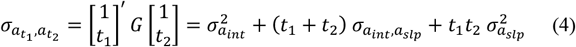

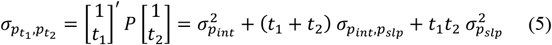

Using these measures of variance and covariance, the heritability of traits at various points and the genetic correlation between points along the gradient *t* were computed as follows:

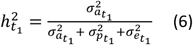

with 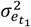 the residual variance of the trait at point *t*_*1*_.

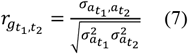

Significance of the different correlations were tested using the method described in [43].

#### Production level at the start of period under study

In particular, equation (6) was applied at *t*_l_=70 weeks of age, to calculate the heritability of the production level at the start of the period under study 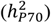.

#### Rate of change of traits between 70 and 90 weeks of age

The average rate of change per week (RC) for a trait X was assessed from 70 weeks of age to the end of the period under study t_max_ (90 weeks of age for RFI and DBWV and 92 weeks of age for the others traits) with the formula

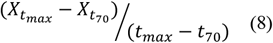

The genetic variance of RC was calculated as in [44]:

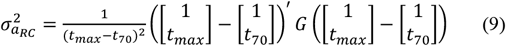

Equation (9) can be simplified as:

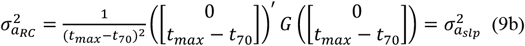

Similarly, the permanent environment variance of OP was calculated as:

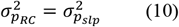

The residual variance of OP was calculated as:

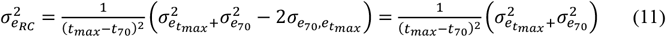

under the same assumption as ASREML of no residual covariance between the phenotypes at 70 weeks of age and t_max_. It can be assumed, however, that if the residual covariance were not zero, it would be positive, as the environmental effects included in the residual would likely affect measurements between 70 and 92 weeks in the same direction. The consequence could be an overestimation of 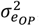,and therefore perhaps a slight underestimation of 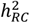 calculated as:

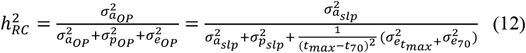

Finally, the genetic correlations between the slope a_slp_ and the intercept a_int_ of the linear function modelling the additive genetic effect was calculated as:

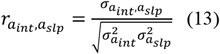

with 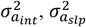 and 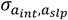 being the genetic (co)variance estimates for the two additive genetic random regression coefficients a_int_ and a_slp_.

### Multivariate analysis

To understand the relationship between the persistence of various traits and pinpoint distinctive clusters among the analysed hens, we used the *FactoMineR* package in R [44] on individual estimated breeding values (EBV) for intercept and slope for the analysed traits (i.e. the random regression coefficient estimates for intercept 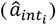 and slope 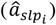). We used the *PCA* function, to identify pattern of association between the EBV for the slope and intercept. To identify distinct persistence clusters, we used the *HCPC* function on the PCA coordinates of each hen persistence. We determined four persistence clusters based on a combination of inertia gain (more than 5% of the total inertia) and the number of animals within each cluster. The significance of the difference between groups has been tested by student tests, with a correction by False Discovery Rate applied to the obtained p values.

## Results

### Population average trajectories through time

The population average trajectories between 70 and 92 weeks of age, calculated as 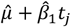 (with 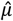 and 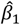 being, respectively the estimates for *μ* and *β*_l_ in model (1)), differed among traits (Figure 2). More specifically, persistence was observed for DBWV, FCR, HU, and YP, as their slopes were close to 0. Increasing trends with age were observed for BW, MEW and ESC as shown the positive slopes observed. On the other hand, significantly negative slopes were observed for the remaining traits

**Fig. 2:**
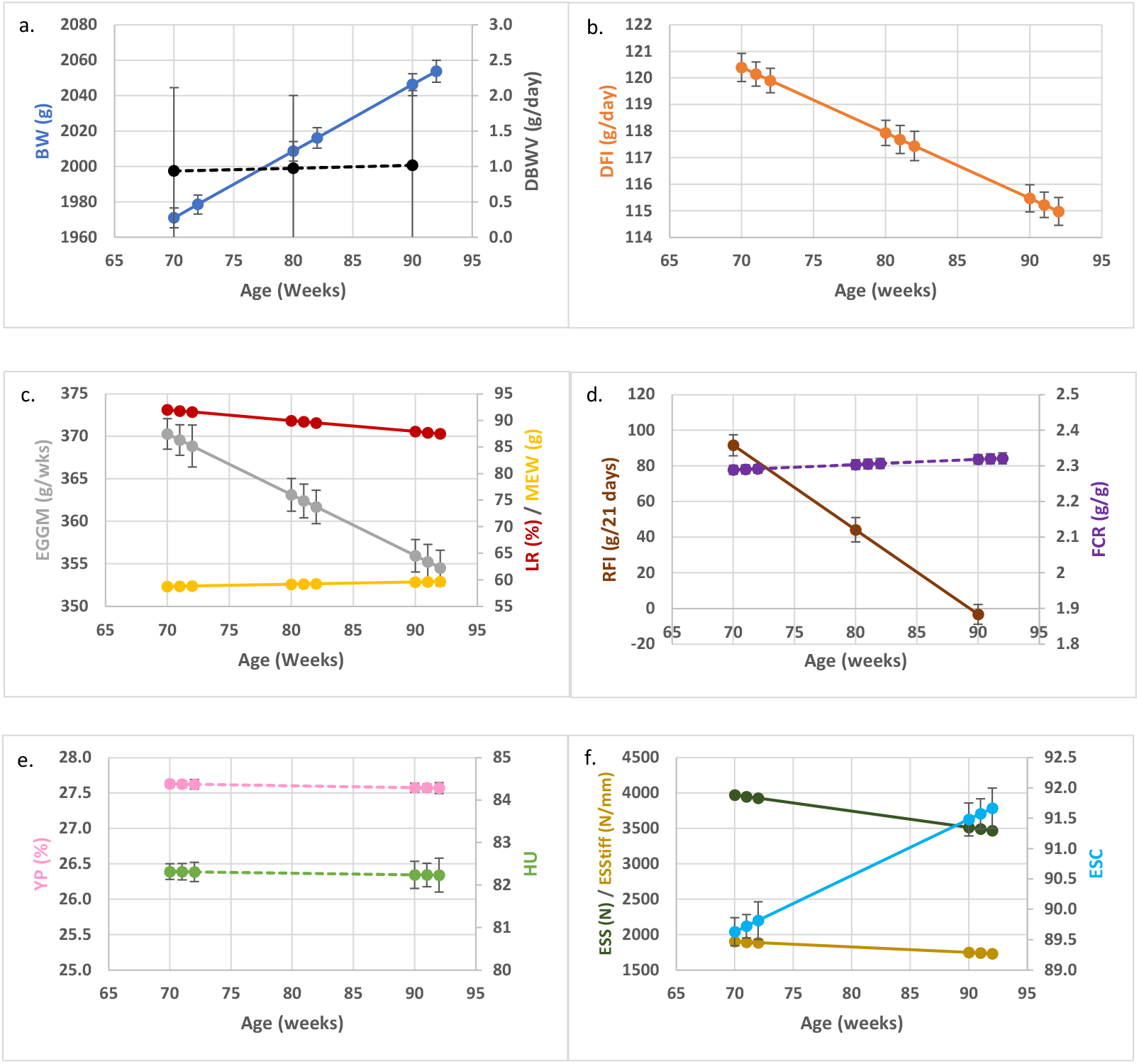
Population average trajectories for BW and DBWV (a), DFI (b), EGGM, EW and LR (c), FCR and RFI (d), HU and YP (e), ESS and ESStiff and ESC (f). Coloured lines represent the mean trait, vertical black lines the standard error. Whole lines represent the traits where the slope is significantly different form 0. Dotted lines represent traits with the slope not significantly different from 0.

### Genetic variance estimates for the intercept and slope of the random regression

The genetic variance estimates of the slope term (σ^2^aslp) were not significantly different from 0 for MEW and ESC indicating limited genetic variability in the rate of change for these traits over time. For the remaining traits, both the intercept and slope showed significantly genetic variance, as detailed in Table 1. For many traits, the genetic covariance between the intercept and the slope was non-significant. Among the traits with significant covariance, a negative covariance between the slope intercept was identified for DBWV, EGGM, and ESS, resulting in negative correlations between these parameters. In contrast, the covariance between the slope and intercept for HU was positive, leading to a positive correlation between them.

**Table 1.** Genetic variance components for the intercept and the slope and heritability estimates for the production at start and rate of change of studied characters.

| Traits | Genetic variance component estimates for the intercept and the slope |  |  |  | Heritabilities |  |
| --- | --- | --- | --- | --- | --- | --- |
| | $\sigma^2_{\text{aint}}^*$ | $\sigma^2_{\text{aslp}}^*$ | $\sigma_{\text{aint,aslp}}^*$ | $r_{\text{aint,aslp}}^*$ | $h^2_{\text{P70}}^*$ | $h^2_{\text{RC}}$ |
| BW | 13810 ± 2842 | 2.95 ± 1.37 | 39.3 ± 46.20<sup>1</sup> | 0.19 ± 0.23<sup>1</sup> | 0.45 ± 0.08 | 0.08 ± 0.04 |
| DBWV | 1.59 ± 0.59 | 0.0136 ± 0.00 | -0.14 ± 0.04 | -0.97 ± 0.04 | 0.12 ± 0.04 | 0.20 ± 0.06 |
| LR | 16 ± 4.60 | 0.035 ± 0.01 | -0.22 ± 0.19<sup>1</sup> | -0.30 ± 0.22<sup>1</sup> | 0.14 ± 0.04 | 0.09 ± 0.03 |
| MEW | 7.0 ± 1.60 | 0.00024 ± 0.00<sup>1</sup> | 0.003 ± 0.024<sup>1</sup> | 0.07 ± 0.61<sup>1</sup> | 0.35 ± 0.07 | 0.01 ± 0.02<sup>1</sup> |
| EGGM | 837 ± 196 | 0.45 ± 0.19 | -13.4 ± 5.30 | -0.69 ± 0.17 | 0.27 ± 0.06 | 0.05 ± 0.02 |
| DFI | 48 ± 13 | 0.16 ± 0.04 | -0.70 ± 0.60<sup>1</sup> | -0.26 ± 0.19<sup>1</sup> | 0.17 ± 0.04 | 0.18 ± 0.04 |
| FCR | 0.012 ± 0.001 | 0.00001 ± 0.00 | -0.0002 ± 0.00<sup>1</sup> | -0.46 ± 0.20<sup>1</sup> | 0.13 ± 0.03 | 0.04 ± 0.02 |
| RFI | 6840 ± 1879 | 45.99 ± 8.25 | 15.4 ± 109.9<sup>1</sup> | 0.03 ± 0.19<sup>1</sup> | 0.27 ± 0.07 | 0.48 ± 0.08 |
| HU | 17.0 ± 3.8 | 0.05 ± 0.01 | 0.30 ± 0.20 | 0.34 ± 0.15 | 0.29 ± 0.06 | 0.23 ± 0.05 |
| YP | 1.60 ± 0.30 | 0.0008 ± 0.00 | -0.004 ± 0.01<sup>1</sup> | -0.11 ± 0.19<sup>1</sup> | 0.41 ± 0.07 | 0.12 ± 0.04 |
| ESS | 99963 ± 28643 | 131.92 ± 45.18 | -2349.5 ± 936.1 | -0.65 ± 0.15 | 0.19 ± 0.05 | 0.09 ± 0.03 |
| ESStiff | 23038 ± 5647 | 10.86 ± 5.19 | -72.3 ± 129<sup>1</sup> | -0.14 ± 0.24<sup>1</sup> | 0.25 ± 0.05 | 0.05 ± 0.02 |
| ESC | 25.40 ± 5.80 | 0.003 ± 0.00<sup>1</sup> | -0.14 ± 0.11<sup>1</sup> | -0.53 ± 0.34<sup>1</sup> | 0.28 ± 0.06 | 0.01 ± 0.02<sup>1</sup> |

<sup>1</sup>: Values with this superscript are not significantly different from 0
\*: $\sigma^2_{\text{aint}}$ and $\sigma^2_{\text{aslp}}$ are the variance estimates for the intercept and slope terms of the additive genetic effect, $\sigma_{\text{aint,aslp}}$ is the covariance, $r_{\text{aint,aslp}}$ is the genetic correlation between the intercept and slope of the additive genetic effect, $h^2_{\text{P70}}$ is the trait heritability at the start of the recording period (70 weeks of age) and the $h^2_{\text{RC}}$ is the heritability of the rate of change.

### Heritability estimates for the production at start and rate of change

Heritability estimates, detailed in Table 1, at the start of the analysed period (P70) were significantly different form 0 for all traits, ranging from 0.12 for DBWV, to 0.45 for BW. Conversely, the heritability of the rate of change (RC) was not significantly different from zero for MEW and ESC. For the other traits it was significant and ranged from 0.05 for ESS_tiff_ to 0.48 for RFI. For most traits, the h^2^_RC_ was smaller than the h^2^_P70_ estimates, while for RFI and DBWV it was notably larger.

### Trait genetic parameter estimation at different ages

Variance component and heritability estimates are represented in Figures 3 to 6 and data are summarised in Supplementary table 2.

**Fig. 3:**
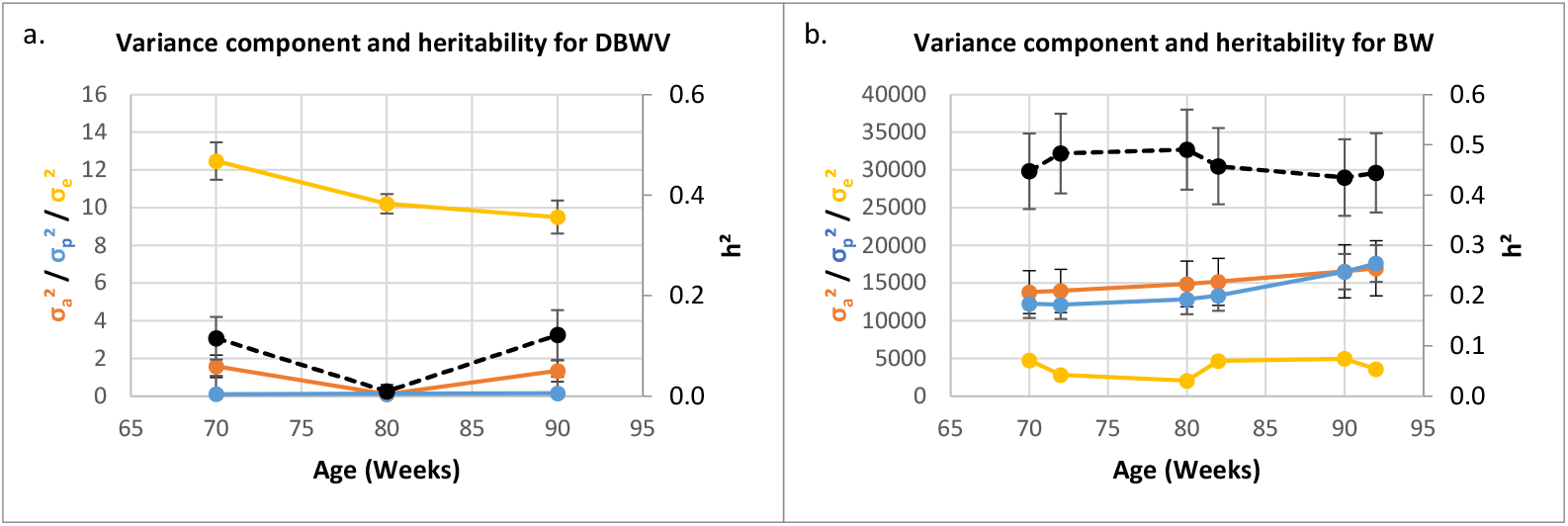
Variance component and heritability for DBWV (a) and BW (b). Orange lines represent the additive genetic variances (σ_a_^2^), blue lines the permanent environment variances (σ_p_^2^), yellow line the residual variances (σ_e_^2^), dotted black line the heritability (h^2^) and vertical black lines the standard error.

For DBWV we found relatively modest heritability at 70 and 90 weeks of age (∼0.12 (± 0.05)). At 80 weeks, the absence of significant genetic variance led to a heritability not significantly different from zero (Figure 3a). The permanent environment variance remained consistently low, while residual variance remained notably high across the entire period. In contrast, for BW a distinct variance pattern emerged, characterised by consistently low residual variance, and increasing genetic and permanent environmental variances over time. The heritability values across ages remained relatively stable, ranging between 0.45 and 0.50 (Figure 3b).

The heritability estimates for LR, were overall low (0.13 ± 0.03) and relatively stable across ages. The permanent environmental variance was higher than the genetic variance, particularly from 80 weeks onwards, and the residual variance was high throughout the period (Figure 4a). For MEW, a low but steady decrease in heritability was observed. Genetic and residual variances were overall stable, while, permanent environment variance increased after 81 weeks of age (Figure 4b). For EGGM a decrease of heritability was observed, with age passing from 0.27 at 70 weeks of age to 0.12 at 92 weeks of age (Figure 4.c). Genetic variance was higher than the permanent environment variance up to 80 weeks of age, when the trend was inverted; the residual variance was high throughout the period.

**Fig. 4:**
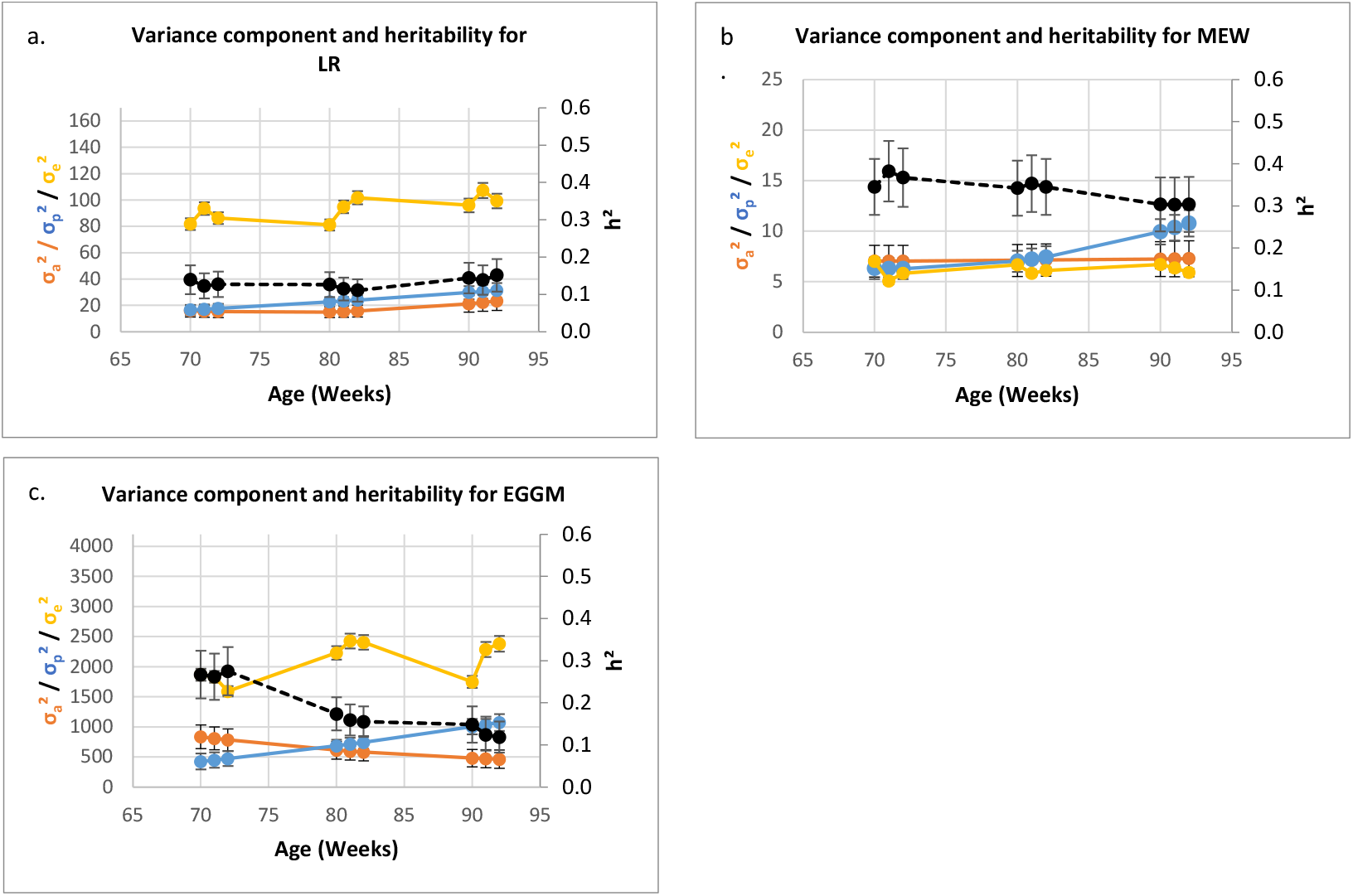
Variance component and heritability for LR (a), MEW (b) and EGGM (c). Orange lines represent the additive genetic variances (σ_a_^2^), blue lines the permanent environment variances (σ_p_^2^), yellow line the residual variances (σ_e_^2^), dotted black line the heritability (h^2^) and vertical black lines the standard error.

Feed-related traits presented high residual variance, particularly over the 80-82 period. The heritability of FCR remained stable with advancing age, being around 0.13 over the period (Figure 5a). Over this period, genetic variance showed a low downward trend, while the permanent environment variance showed an upward trend. The heritability of DFI remained stable between 71 and 80 weeks, dropped between 81 and 82 weeks, for then increase during the 90-92 weeks period. In contrast, residual variance followed an opposite pattern, peaking between 81 and 82 weeks of age before declining in the 90-92 weeks period. Both genetic and the permanent environment variances increased from 82 weeks onwards.

**Fig. 5:**
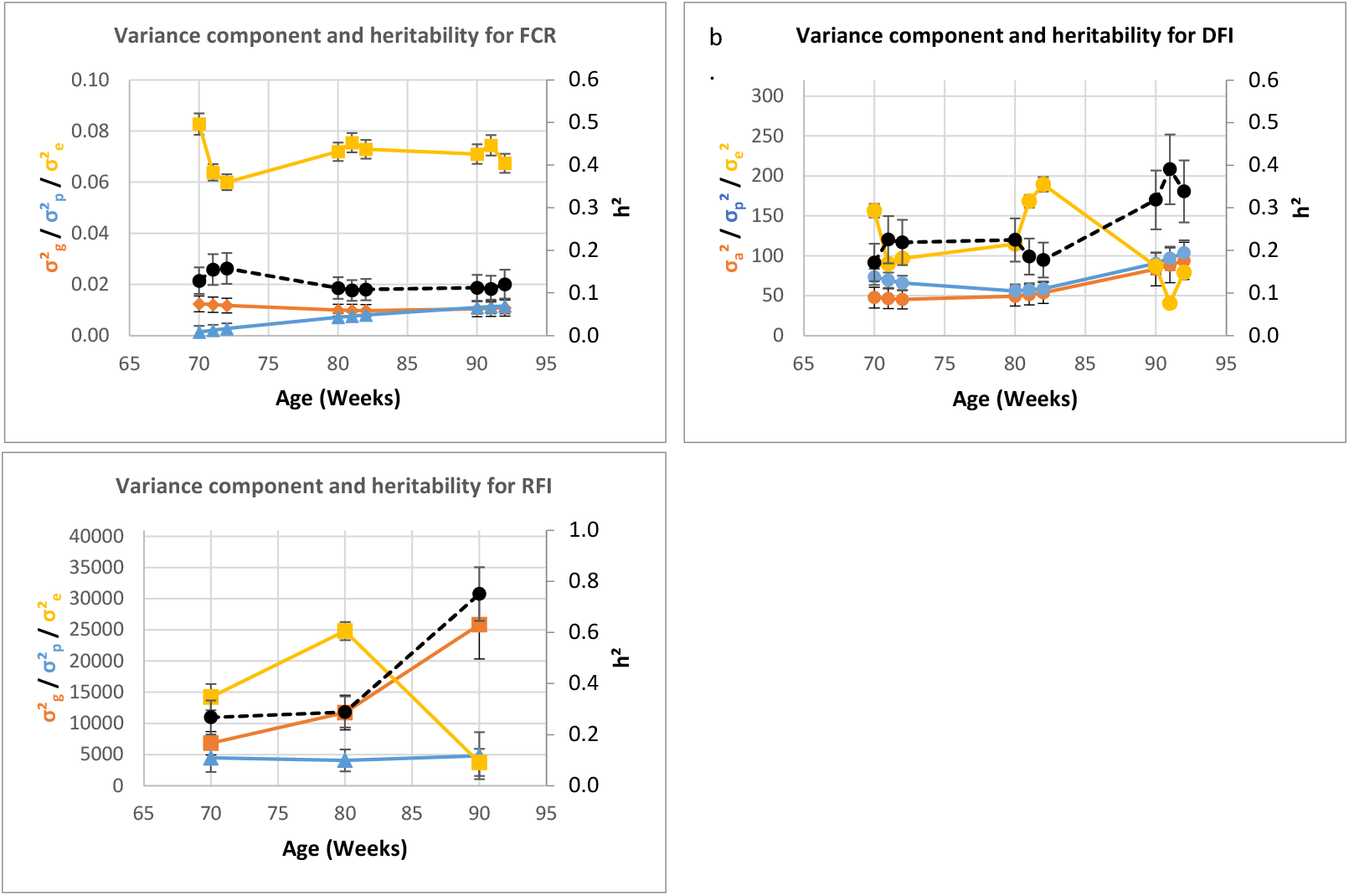
Variance component and heritability for FCR (a), DFI (b) and RFI (c). Orange lines represent the additive genetic variances (σ_a_^2^), blue lines the permanent environment variances (σ_p_^2^), yellow line the residual variances (σ_e_^2^), dotted black line the heritability (h^2^) and vertical black lines the standard error.

For the RFI, the genetic variance increased from 70 to 90 weeks, while the permanent environment one remained stable. The residual variance for this trait showed the same patterns of the one for DFI, an increase from 70 to 80 weeks, followed by a decrease to 90 weeks. Heritability remained stable until 80 weeks, around 0.27. After that a strong increase could be observed (Figure 5c).

In terms of egg quality-related traits, both HU and YP exhibited an overall increase in heritability with age, and a genetic variance that was higher than the permanent environment and residual variances at 90 weeks of age (Figure 6a and Figure 6b). Eggshell quality traits presented similar pattern of variance components, characterized by elevated residual variances and relatively low genetic variances. For ESC and ESS the heritability declined with age (Figure 6c and 6e), whereas for ESStiff, it remained relatively constant (Figure 6d).

**Fig. 6:**
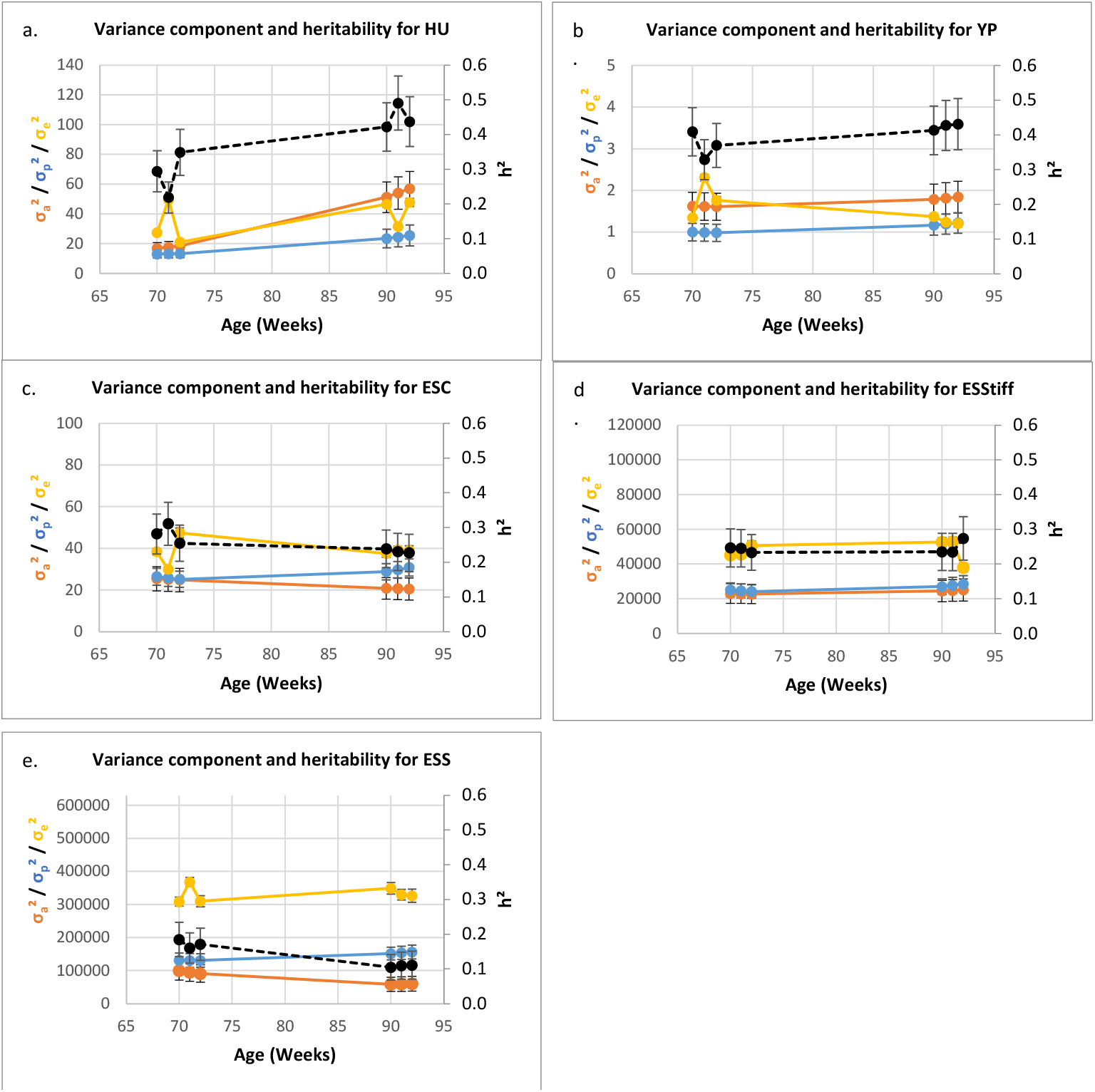
Variance component and heritability for HU (a), YP (b). ESC (c), ESStiff (d) and ESS (e). Orange lines represent the additive genetic variances (σ_a_^2^), blue lines the permanent environment variances (σ_p_^2^), yellow line the residual variances (σ_e_^2^), dotted black line the heritability (h^2^) and vertical black lines the standard error.

Most genetic correlations for traits recorded at one-week intervals within each of the 3 recording periods were close to 1 (supplementary Figure 1). Therefore, in Table 2, we report only the genetic correlations between traits recorded at 70, 80 and 90 weeks of age. For most traits, we observed a good degree of consistency in genetic effects between ages. This was not the case for DBWV that presented genetic correlations not significantly different from 0 in the comparison 70×80 and 80×90 and a strong negative genetic correlation in the comparison 70×90. A reduced genetic correlation at the 70 x 90 wks comparison is observed for LR, DFI, RFI and ESS.

**Table 2.** Genetic correlations (± SE) between traits measured at different ages.

| Traits | 70x80 | 80x90 | 70x90 |
| --- | --- | --- | --- |
| BW | 0.99 ± 0.01 | 0.99 ± 0.01 | 0.97 ± 0.02 |
| DBWV | 0.41 ± 0.45<sup>1</sup> | 0.11 ± 0.57<sup>1</sup> | -0.86 ± 0.18 |
| LR | 0.89 ± 0.04 | 0.92 ± 0.04 | 0.63 ± 0.13 |
| MEW | 1.00 ± 0.00 | 1.00 ± 0.00 | 0.99 ± 0.02 |
| EGGM | 0.98 ± 0.01 | 0.97 ± 0.02 | 0.90 ± 0.06 |
| DFI | 0.84 ± 0.05 | 0.91 ± 0.03 | 0.53 ± 0.13 |
| FCR | 0.95 ± 0.03 | 0.93 ± 0.04 | 0.77 ± 0.13 |
| RFI | 0.78 ± 0.07 | 0.95 ± 0.02 | 0.54 ± 0.13 |
| HU | 0.91 ± 0.03 | 0.97 ± 0.01 | 0.79 ± 0.06 |
| YP | 0.97 ± 0.01 | 0.98 ± 0.01 | 0.91 ± 0.03 |
| ESS | 0.94 ± 0.03 | 0.9 ± 0.05 | 0.69 ± 0.13 |
| ESSstiff | 0.98 ± 0.01 | 0.98 ± 0.01 | 0.91 ± 0.05 |
| ESC | 1.00 ± 0.01 | 0.99 ± 0.01 | 0.98 ± 0.03 |

- : Values not different from 0

**Table 3.** Mean (± SE) estimated breeding values of the slope and intercept for the analysed traits, calculated from each of the 4 cluster groups identified by HPCP analysis.

| Traits | EBV | Cluster 1 | Cluster 2 | Cluster 3 | Cluster 4 |
| --- | --- | --- | --- | --- | --- |
| BW | Intercept (g) | -29.228 ± 5.535<sup>b</sup> | 35.048 ± 5.348<sup>d</sup> | 12.52 ± 4.545<sup>c</sup> | -42.719 ± 6.598<sup>a</sup> |
| | Slope (g/wks) | - | 0.609 ± 0.049<sup>c</sup> | -0.289 ± 0.035<sup>b</sup> | -0.537 ± 0.074<sup>a</sup> |
| D<sup>B</sup>WV | Intercept (g/day) | -6.204 ± 0.259<sup>a</sup> | -4.744 ± 0.343<sup>b</sup> | 6.217 ± 0.309<sup>d</sup> | 4.417 ± 0.58<sup>c</sup> |
| | Slope (g/day/wks) | 0.5638 ± 0.0245<sup>d</sup> | 04696 ± 0.0325<sup>c</sup> | -0.5951 ± 0.0286<sup>a</sup> | -0.3963 ± 0.056<sup>b</sup> |
| LR | Intercept (%) | 1.814 ± 0.114<sup>d</sup> | -0.792 ± 0.141<sup>b</sup> | 0.559 ± 0.094<sup>c</sup> | -3.294 ± 0.176<sup>a</sup> |
| | Slope (%/wks) | -0.106 ± 0.004<sup>a</sup> | -0.054 ± 0.005<sup>b</sup> | 0.072 ± 0.005<sup>c</sup> | 0.112 ± 0.008<sup>d</sup> |
| MEW | Intercept (g) | 0.338 ± 0.11<sup>b</sup> | -0.469 ± 0.125<sup>a</sup> | 0.254 ± 0.097<sup>b</sup> | -0.398 ± 0.178<sup>a</sup> |
|  | Slope (g/wks) | - | - | - | - |
| EGGM | Intercept (g/wks) | 16.993 ± 0.855<sup>d</sup> | -4.599 ± 1.086<sup>b</sup> | 1.469 ± 0.748<sup>c</sup> | -26.506 ± 1.599<sup>a</sup> |
| | Slope (g/wks<sup>2</sup>) | -0.391 ± 0.013<sup>a</sup> | -0.061 ± 0.016<sup>b</sup> | 0.138 ± 0.014<sup>c</sup> | 0.481 ± 0.032<sup>d</sup> |
| DFI | Intercept (g/day) | - | 1.603 ± 0.295<sup>b</sup> | - | -1.895 ± 0.357<sup>a</sup> |
| | Slope (g/day/wks) | -0.159 ± 0.01<sup>a</sup> | -0.173 ± 0.011<sup>a</sup> | 0.195 ± 0.013<sup>c</sup> | 0.117 ± 0.025<sup>b</sup> |
| FCR | Intercept (g/g) | -0.053 ± 0.003<sup>a</sup> | 0.069 ± 0.004<sup>d</sup> | -0.034 ± 0.003<sup>b</sup> | 0.057 ± 0.005<sup>c</sup> |
| | Slope (g/g/wks) | - | -0.0019 ± 0.0001<sup>a</sup> | 0.0013 ± 0.0001<sup>b</sup> | - |
| RFI | Intercept (g/21/days) | -20.992 ± 2.943<sup>a</sup> | 27.471 ± 3.105<sup>b</sup> | -18.057 ± 2.631<sup>b</sup> | 33.671 ± 3.511<sup>a</sup> |
| | Slope (g/21 days/wks) | -4.454 ± 0.235<sup>b</sup> | -5.423 ± 0.2355<sup>a</sup> | 5.2255 ± 0.229<sup>d</sup> | 0.44467 ± 0.361<sup>c</sup> |
| HU | Intercept | -1.151 ± 0.2<sup>a</sup> | 0.57 ± 0.207<sup>b</sup> | - | 1.551 ± 0.201<sup>c</sup> |
| | Slope (/weeks) | -0.082 ± 0.012<sup>a</sup> | 0.024 ± 0.009<sup>b</sup> | 0.008 ± 0.008<sup>b</sup> | 0.086 ± 0.011<sup>c</sup> |
| YP | Intercept (%) | -0.483 ± 0.054<sup>a</sup> | 0.491 ± 0.052<sup>c</sup> | - | 0.206 ± 0.082<sup>b</sup> |
| | Slope (%/wks) | -0.012 ± 0.001<sup>a</sup> | -0.006 ± 0.001<sup>b</sup> | 0.008 ± 0.001<sup>c</sup> | 0.014 ± 0.001<sup>d</sup> |
| ESS | Intercept (N) | - | 115.207 ± 10.905<sup>c</sup> | -111.872 ± 8.008<sup>a</sup> | 63.558 ± 17.744<sup>b</sup> |
| | Slope (N/wks) | -3.112 ± 0.355<sup>a</sup> | -3.793 ± 0.359<sup>a</sup> | 5.313 ± 0.253<sup>b</sup> | - |
| ESSStiff | Intercept (N/mm) | -35.928 ± 6.011<sup>a</sup> | 60.407 ± 6.202<sup>c</sup> | -18.484 ± 4.975<sup>b</sup> | - |
| | Slope (N/mm/wks) | - | -0.332 ± 0.082<sup>c</sup> | 0.47 ± 0.083<sup>b</sup> | -0.776 ± 0.15<sup>a</sup> |
| ESC | Intercept | -0.835 ± 0.226<sup>a</sup> | 1.342 ± 0.211<sup>b</sup> | - | -0.985 ± 0.318<sup>a</sup> |
|  | Slope (/wks) | - | - | - | - |
| Number of animals |  | 247 | 254 | 255 | 350 |

a,b,c,d : Within a line, values with same superscript are not significantly different
-: Value not different from 0

### Principal component analysis and hierarchical clustering of individual breeding value estimated from the trait intercept and slope

The principal component was performed using the EBV estimated for each trait’s intercept and slope but for clarity reasons, only traits having the strongest contribution to the principal components (PC) (cos^2^ value above 0.5 with one of the different PC) were shown in Figure 7.

**Fig. 7:**
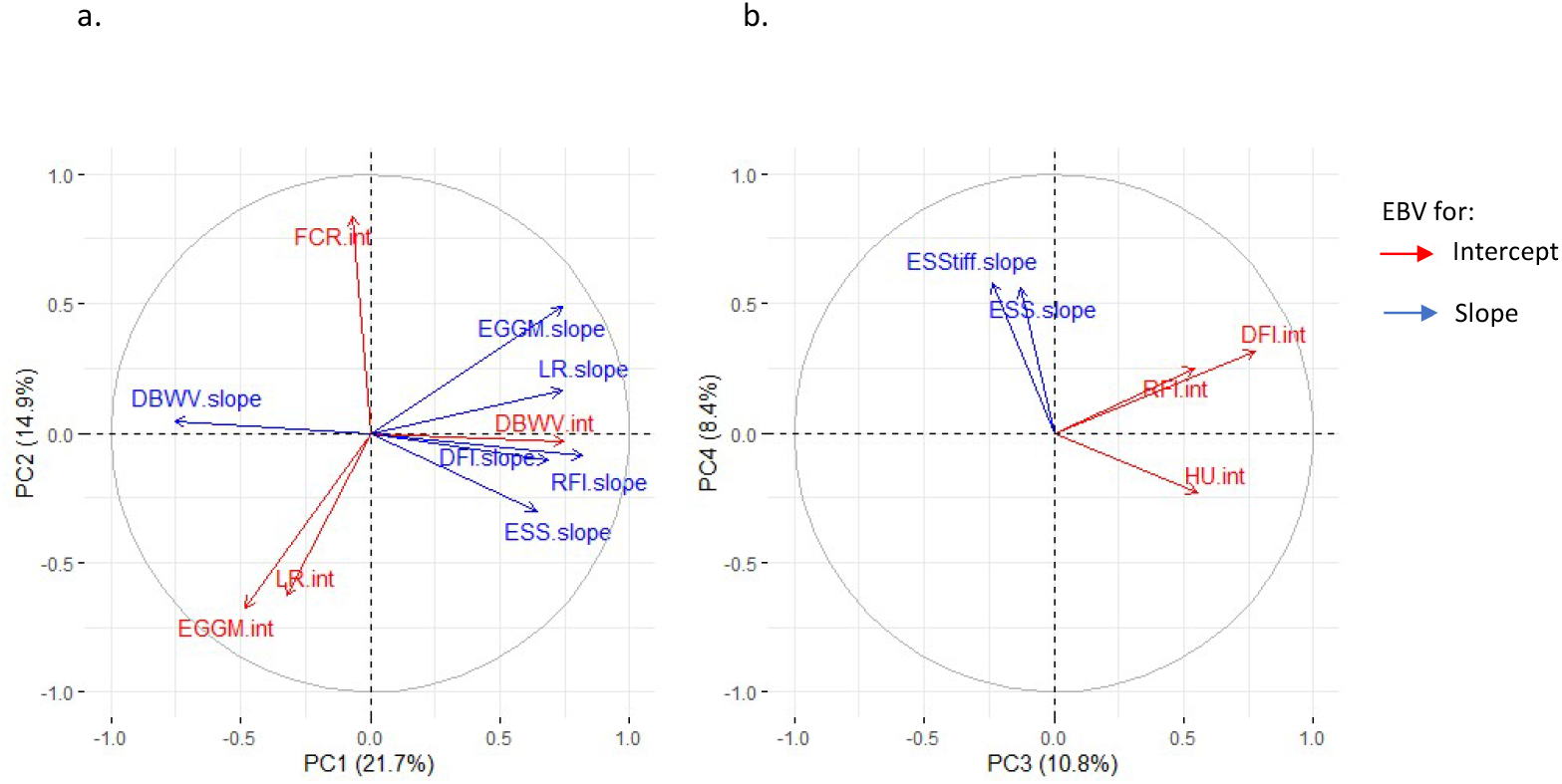
Correlation circles of PC1 and 2 (a) and of PC3 and 4 (b). Blue and red arrows represent the EBV of the slope and intercept of the traits, respectively.

The first PC captured 21.6% of the variability and separated animals based on the EBV for the intercept and slope of DBWV, and the slopes of LR, EGGM, DFI, RFI and ESS. The second PC contributed to 14.9% of the variance and separated animals based on the EBV of the intercept of FCR, LR, and EGGM.

The third PC explained 10.8% of the variance, and separated animals based on the EBV of the intercept of HU, DFI and RFI. Finally, the fourth PC, explained 8.4% of the variance and separated animals based on the slope of ESS and ESStiff.

The hierarchical clustering analysis allowed to identify four distinct clusters: cluster 1 included 254 animals, cluster 2 included 255 animals, cluster 3 included 350 animals, and cluster 4 included 138 animals. The cluster dendrogram is presented in Figure 8a and the cluster mean EBV values for the intercept and slope of each trait are provided in the Table 3. The projection of the four clusters on the first and second PC showed that the first axis primarily separated cluster 1 and 2 from the cluster 3 and 4, while the second mainly separated cluster 1 and 3 from clusters 2 and 4 (Figure 8b). The subsequent PC axes did not provide clear separation between clusters and were consequently not shown.

**Fig. 8:**
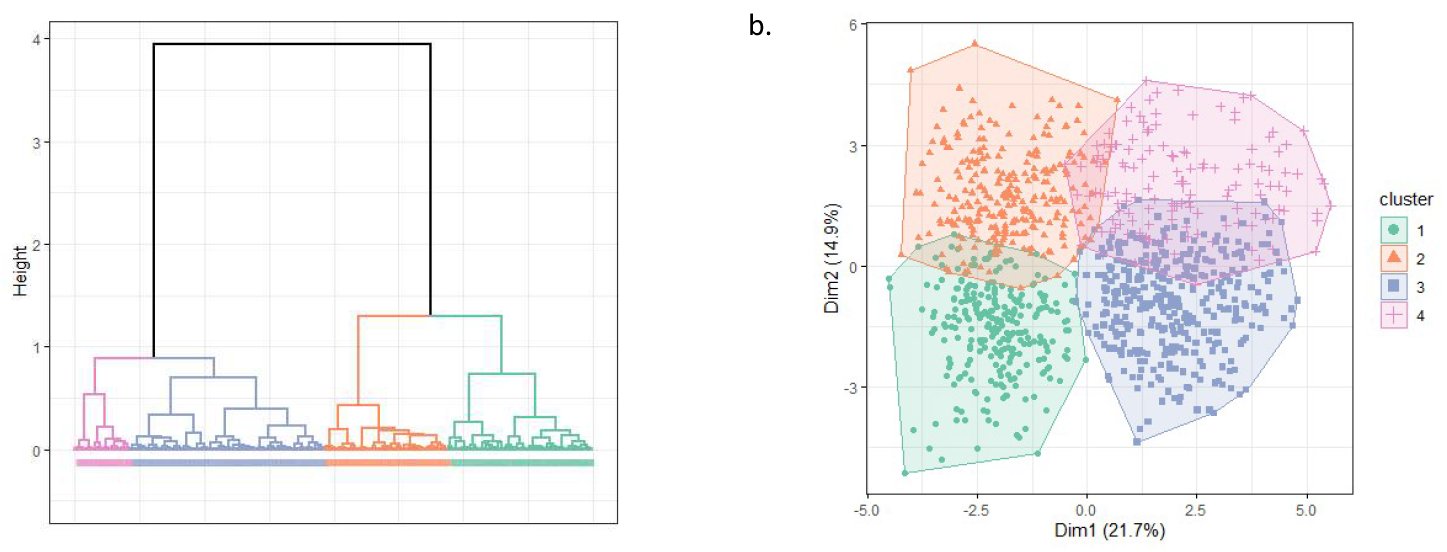
Hierarchical clustering (a) and projection of the different clusters on the first and second principal components (b)

The principal component separated animals primarily based on the EBV trajectories of the traits, while the second component separated animals based on the EBVs of traits at 70 weeks of age. Hens from clusters 3 and 4 contrast with those from cluster 1 and 2 (separation based on PC1) by showing improving EBVs for the slope of LR over time, as well as their positive EBVs for the slopes of DFI and RFI. Conversely, hens from clusters 1 and 3 differ from those from clusters 2 and 4 (separation based on PC2) by having negative EBVs for MEW and positive EBV for egg shell quality traits at 70 weeks of age (Table 3 and supplementary figure 2).

## Discussion

In this study, we investigated the persistence of traits related to egg production, egg quality, body weight, and feed efficiency in laying hens at advanced production stages (from 70 to 92 weeks of age). Persistence is defined as the ability to sustain a trait’s performance over time. Our goal was to better understand the genetic factors influencing hens’ ability to sustain performance at advanced production ages and to identify opportunities for selecting hens with improved persistence.

### Phenotypic evolution as a function of age

The observed decrease in laying rate with age aligns with previous studies, including those that analysed laying rate at different ages (raw data) [16, 18, 45] as well as studies that, like our, employed random regression techniques [20, 46–48]. Similarly, the increase in egg weight at later production stages, which is expected in aging hens and has been described previously [2, 45, 49], was also observed in our study. This increase observed from 70 to 93 week of age is nevertheless rather limited (1.6%), reflecting the successful selection strategy on egg weight stability applied on this parental line.

Stability in egg quality is also a major breeding goal in egg production especially in the context of extending the production period [6]. In our study, different aging trajectories were observed for egg quality traits: the internal egg quality traits remained stable, while shell quality traits declined over time. The persistence of HU values in our study contrasts with previous findings, which reported a general decline in albumen quality in aging hens [2, 50, 51]. This stability is likely the result of the selection strategies in commercial flocks aimed at maintaining stable egg weight and improving albumen quality [3]. For the shell quality traits we observed a loss of resistance with age, aligning with the well-documented increase in eggshell fragility in aging hens [52, 53]. Unfortunately, the absence of shell weight and mineral composition measures in our dataset prevents us from verifying whether the increased fragility of the shell stems from a decreased shell proportion, diminished calcium composition, or alterations in the shell’s structural integrity [53].

Most studies that investigated persistence in hens, mainly focused on laying rate or egg quality [54, 55]. In this study, we explored the persistence of a broader range of traits, including feed intake and feed efficiency related traits, which to our knowledge, have never been investigated for ages beyond 70 weeks of age. FCR and RFI followed different trajectories: FCR showed a steady trend, while RFI decreased over time. This difference can be justified considering that these two metrics give different perspectives on the animal’s efficiency, with FCR being a gross measure of feed intake relative to production, that does not consider maintenance requirements as instead does RFI [56]. Indeed, the fact that both EGGM and DFI account for the same weight over the FCR explain its steadiness, the decrease in both of these traits being very close. Conversely, the improvement in RFI observed in aging hens, may be attributed to the small increase in BW despite a steady reduction in DFI and EGGM, which could reflect a reduction of maintenance requirements with age.

Despite a decline in feed intake throughout the period, we observed a gain in BW. Indeed, increase in body weight is often attributed to the accumulation of abdominal fat, which is linked to physiological factors mirroring metabolic and hormonal changes [57]. Increased abdominal fat was also observed in our study, where a larger abdominal fat proportion was observed in birds at 90 weeks of age compared to those analysed at 70 weeks (2.4% versus 3.4%, p <0.05, data not shown).

### Kinetics of genetic architecture of traits

For many traits, heritability estimates were modest across ages, remaining relatively stable for some, while fluctuating for others, with values often falling below expectations based on earlier reports. This is notably the case for LR and EGGM where heritability estimates at all age points were lower than those reported in other studies at earlier production ages [3, 58–62], suggesting that the genetic contribution to these traits diminishes over time, particularly in later production stages.

Residual variances were particularly high for egg production traits, FCR and eggshell quality traits. This highlights the substantial role of non-genetic factors for these traits too, suggesting that short terms fluctuations or physiological changes due to aging may dominate phenotypic expression of these traits during advanced production life stages. For egg production, this may be a reflection of the intense selection applied to this trait before 70 weeks of age [6] that may lead to an erosion of the available genetic variability at later ages, with a consequent increased role of non-genetic factors in egg production variation.

Notably, we observe a large increase in residual variance for DFI and feed efficient related traits at 80-82 weeks of age, which may have distinct origins. The extended production period might require physiological adaptations, and the 80–82 weeks of age period could mark a critical step in the animal’s energy requirements and allocation. This turning point could explain the reduced genetic correlations between these traits measured at 70 and 90 weeks of age compared those observed between 70 and 80 weeks of age. Alternatively, temporary stress might explain the observed fluctuations in feed intake and efficiency. This stress may have resulted from the transition from collective feeders (used by multiple hens) to individual feeders (designated for individual use) a week before the beginning of the three recording periods (i.e., at 70, 80 and 90 weeks of age). We cannot exclude that for unforeseen reasons, the change of feeders for the measure at 80-82 weeks of age might have been more stressful compared the other recording periods.

The high residual variance resulted in a scattered pattern of heritability estimates for DFI and RFI across the three recording periods. Heritability estimates for DFI in the literature vary considerably across studies and genotypes, ranging from 0.20-0.26 in a reciprocal cross between White Leghorns and Dongxiang blue-shelled chickens cross [63], in line with the values we observed in the first two measured periods, to 0.46 in an experimental brown-egg pure line layer population [64], which aligns with our findings at 90 weeks of age. The RFI heritability estimate at 70 and 80 week of age were in the range of what proposed earlier on different laying population [37, 63–66]. Conversely, the estimates at 90 weeks were out of the range found for this character, being even higher with what observed in broilers [67]. It is interesting to note that the scattered heritability and residual variance values across ages obtained in our study highlight the high sensitivity of DFI to environmental changes and stressors compared to other traits. Animals typically respond to perturbations by rapidly reducing their feed intake and possibly recovering once the stressor ends. In pigs, the variability of daily feed intake is considered a particularly promising metric for the development of resilience indicators [68] which can be used as a clinical sign of disease [69]. Further studies will be required to validate this in laying chickens.

Finally, contrasted heritability patterns were also observed among egg quality traits. Those related to shell quality were primarily influenced by high residual variance, whereas internal egg quality traits were affected by additive genetic and residual variances. Heritability values for shell quality traits were at the lower end of the spectrum reported in the literature, ranging from 0.10 to 0.48 for ESS, 0.2 to 0.49 for ESStiff, and 0.17 to 0.46 for ESC, depending of the strain and period of measure [62, 70–74]. The low heritability estimates, coupled with high residual variances, suggest that the genetic control of eggshell strength becomes less influential with advancing age, making quality more reliant on other non-genetic factors. This is further supported by the reduced genetic correlation for ESS in the comparison between 70 and 90 weeks of age, indicating a loss of genetic control over trait variation with age. Among the non-genetic factors contributing to variability in eggshell quality of older laying hens, are the health and aging of endometrial cells that lead to a reduced calcium-transporting ability [75].

Previous studies have reported that albumen quality typically decreases with advancing production ages, as evidenced by reduced Haugh unit scores and increased variability [76, 77]. However, our results do not show such a deterioration. On the contrary, HU estimates exhibit good persistence, and relatively high heritability, particularly at 90 weeks of age. This is accompanied by a growing genetic variance, which triples from 70 to 90 weeks, highlighting opportunities to capitalize on genetic potential to maintain acceptable albumen quality over longer production periods.

### Genetic Architecture of Trait Stability Over Time

Genetic improvement at 70 weeks of age is indeed feasible for most traits, given the relatively high heritability estimates, which suggest that genetic determinism remains strong at advanced production stages for most analysed traits. In contrast, heritability estimates for the rate of change over time were very low for most traits, including egg production traits. As a result, any attempt to modify it through genetic selection, whether to increase it or reduce it, will be slow or less effective, as non-genetic factors play a major role in the variance of a trait’s rate of change over time, as already found in other species [78–80].

The estimation of the genetic correlation between the intercept and the slope for each of the traits over the studied interval, helps understand how genetic improvement of a trait at 70 weeks of age influences its overall evolution at later ages. The fact that most traits present genetic correlation between the intercept and the slope that are not significantly different from zero suggests that improving a trait at 70 weeks is unlikely to affect its rate of change at later production stages relative to the population average. This because the genetic factors influencing an individual’s baseline performance (estimated at 70 weeks of age in this study) do not affect the persistence or trajectory of the trait over time. This has already been demonstrated in pigs, where QTL regions associated with the slope are fewer and do not overlap with those linked to the intercept for both growth and feed intake traits [81], underlying the fact that different genetic mechanisms must be involved in the control of the performance at a given age compared to its long-term stability.

Potential trade-offs between initial value and long-term trend may be expected considering the significant negative genetic correlations between the intercept and the rate of change observed for DBWV, EGGM and ESS. The expected consequence is that a genetic improvement at 70 weeks will lead to a decline of the trait values over time, requiring careful management in breeding programs. HU presented instead a positive genetic correlation indicating that an improvement of this strait at earlier ages can have long-term benefits.

### Trait Dynamics Based on EBV Clustering

The PCA provided insight into the relationship between the EBVs of the slope and intercept of analysed traits. The correlation circle of the PCA confirmed the genetic correlation observed between the slope and intercept for DBWV, EGGM and HU. Interesting, the relationship between the EBV estimation of DBWV intercept and that for LR and EGGM slopes, highlights the importance of DBWV at 70 weeks of age, in determining the trajectory of egg production at advanced ages. Hens with above average EBV for DBWV at 70 weeks of age had also a higher potential to maintain laying rate over time. It is suggested that leaner hens may be more susceptible for nutritional deficits than heavier ones [82]. Indeed, previous studies have shown that heavier hens tend to have a higher egg production compared to light hens despite equal feeding conditions during rearing [83]. This is in agreement with the observations that body weight of laying hens not only influences early-stage egg production but also plays an important role in the hen’s ability to sustain production at later stages [84, 85].

The EBV of DBWV intercept was also positively correlated with the EBV of RFI and DFI slopes, indicating that hens with larger body gains at 70 weeks of age tend to be less efficient in the long term, mostly due to an increase in feed intake as previously observed [63]. This reinforces the awareness that genetic selection for egg production persistence must address feed efficiency too, to avoid selecting hens that maintain high productivity but with excessive feed intake, increasing by consequence production costs and environmental impact [5, 86].

In the parental line analysed in this study, the clustering analysis reveals distinct genetic profiles in terms of both egg production and quality, as well as body weight and feed efficiency dynamics. If the breeding goal is to improve egg persistence at advanced production stages, then animals from Cluster 3 can be good candidates for selection. Hens from this cluster presented positive EBV for LR, at 70 weeks of age and demonstrated a good genetic potential for egg persistence at advanced stages of production. It is interesting to note that these same hens also showed rather stable BW despite an increased in DFI as the hens aged. This pattern could reflect an ability to maintain energy allocation toward egg production, rather than body weight gain, offering benefits for long-term productivity. The counterpart is that their genetic potential for increased feed consumption and by consequence, decreased feed efficiency over time could represent a challenge, especially in commercial settings where feed costs are a major factor of egg cost production. The EBV patterns for LR, DFI, and RFI observe across the four Clusters suggest the existence of a trade-off among traits, where lower feed intake may limit hens’ egg production but improve feed efficiency.

The relationship between egg weight and shell quality is also evident across clusters. Hens from Clusters 1 and 3 presented positive EBVs for egg weight at 70 weeks, while those form clusters 2 and 4 presented negative EBVs. Interestingly, the positive EBVs for egg weight in Clusters 1 and 3 is accompanied by negative EBVs for eggshell quality, which aligns with the known negative relationships between egg size and shell strength [87]. In contrast, clusters 2 and 4, which have negative EBVs for egg weight, show positive EBVs for eggshell quality traits, confirming the known tendency for smaller eggs with stronger shells.

These results highlight the complex relationships between laying performance, feed efficiency, body weight stability, and egg quality. While clusters 3 and 4 demonstrate the ability to sustain or even improve egg production, they face challenges in feed efficiency and body weight fluctuations. On the other hand, clusters 1 and 2, with better feed efficiency and potentially more stable body weight, may experience limitations in long-term egg production and egg quality.

## Conclusion

The current study aimed to gain a better understanding of the persistence capacity of hens for a broad range of traits, including egg production, quality and feed efficiency related traits, at advanced production periods, from 70 to 92 weeks of age. The estimation of genetic parameter at different ages revealed high genetic correlations between traits across different ages, indicating consistent genetic influence on a given trait across different life times. Selection to genetically improve egg production persistence may be possible, but it may be challenging due to the low genetic variances. Moreover, the existence of distinctive response patterns associated with extending the production period, underlies the existence of compromises among traits therefore selecting for enhanced persistence across multiple traits will inevitably require making compromising with respect to breeding goals.

## Abbreviations

a*: Redness of the eggshell in chromameter readings
a_int_: Random additive genetic effect for the intercept of the random regression
a_slp_: Random additive genetic effect for the regression coefficient for slope of the random regression
b*: Yellowness of the eggshell in chromameter readings
β_1_: Fixed regression coefficient on the time period between the day of egg lay and the day of quality measurement
β_2_: Fixed regression coefficient for the slope on the age in weeks
BW: Body Weight
DBWV: Daily Body Weight Variation
DFI: Daily Feed Intake
DOB: Date of Birth
EBV: Estimated Breeding Values
EggLaid: Number of Eggs Laid
EGGM: Egg Mass
e_ijkl_: Random residual effect
ENVT55wk: Rearing system before 55 weeks
ESC: Eggshell Color
ESS: Eggshell Breaking Strength
ESStiff: Eggshell Stiffness
EW: Egg Weight
FCR: Feed Conversion Ratio
FI: Feed Intake
HCPC: Hierarchical Clustering on Principal Components
HU: Haugh Unit
L*: Lightness of the eggshell in chromameter readings
LR: Laying Rate
MEW: Mean Egg Weight
P70: Production level at start
p_int_: Random permanent environment effect for the intercept of the random regression
p_slp_: Random permanent environment effect for the regression coefficient for slope of the random regression
PC: Principal Component
PCA: Principal Component Analysis
PFI: Predicted Feed Intake
PosCage: Individual cage location in the barn during the period of measure
RC: Rate of Change
REML: Restricted Maximum Likelihood
RFI: Residual Feed Intake
RRM: Random Regression Models
TimeRM: Time between the day of egg lay and the day of quality measurement
WFI: Weekly Feed Intake
YP: Yolk Percentage

## Acknowledgements

We thank Novogen for producing and rearing the animals and for sharing with us phenotypes and genealogy data and XXX for the English editing services.

## Authors ‘contributions

TB, SL, and TZ conceived the project and secured funding and QB, NB, TZ, TT and PL refined the project and advised on statistical analyses. TB managed animal production and organised phenotype collection. QB performed the statistical analysis and QB and TZ wrote the manuscript. NB, and TT contributed useful comments and suggestions on the manuscript draft. All authors read and approved the final manuscript.

## Availability of data and materials

The datasets used in the study are available from the corresponding author on request.

## Ethics approval and consent to participate Consent for publication

Not applicable

## Figure titles

**Supplementary table 1:** Variance estimates for the intercept (σ_p_^2^_int_) and slope terms (σ_p_^2^_slp_) of the permanent environment effect, the covariance (σ_pint, pslp_), the correlation between the intercept and slope of the permanent environment effect (r_pint,pslp_).

| Traits | Variance component estimates for the intercept and the slope |  |  |  |
| --- | --- | --- | --- | --- |
| | $\sigma_{p^2_{int}}$ | $\sigma_{p^2_{slp}}$ | $\sigma_{pint,slp}$ | $r_{pInt,slp}$ |
| BW | 12256 ± 1865 | 15.43 ± 1.6 | -47.9 ± 37.8 | -0.11 ± 0.09 |
| DBWV | 11.38 ± 0.95 | 0.0009 ± 0.00001 | 0.1 ± 0.05 | 0.96 ± 1.0 |
| LR | 16.6 ± 3.9 | 0.006 ± 0.012 | 0.27 ± 0.18 | 0.86 ± 1.0 |
| MEW | 6.3 ± 1.1 | 0.01108 ± 0.0012 | -0.021 ± 0.024 | -0.08 ± 0.09 |
| EGGM | 428 ± 133 | 0.3 ± 0.25 | 11.4 ± 4.4 | 1.0 ± 0.78 |
| DFI | 74 ± 10 | 0.26 ± 0.03 | -2.2 ± 0.5 | -0.50 ± 0.08 |
| FCR | 0.0014 ± 0.0024 | 0.00009 ± 0.00001 | 0.0003 ± 0.0001 | 0.91 ± 0.94 |
| RFI | 4477 ± 2261 | 5.65 ± 10.9 | -47.8 ± 12.5 | -0.30 ± 0.25 |
| HU | 13.1 ± 3.0 | 0.02 ± 0.01 | 0.03 ± 0.10 | 0.05 ± 0.19 |
| YP | 1.0 ± 0.2 | 0.0009 ± 0.0002 | -0.005 ± 0.005 | -0.16 ± 0.16 |
| ESS | 131346 ± 21818 | 67.21 ± 41.49 | -166.4 ± 756.6 | -0.06 ± 0.24 |
| ESStiff | 25123 ± 4065 | 31.53 ± 6.42 | -267.6 ± 119.5 | -0.30 ± 0.11 |
| ESC | 26.4 ± 4.1 | 0.04 ± 0.005 | -0.34 ± 0.10 | -0.33 ± 0.09 |

**Supplementary table 2:** Variance and heritability estimates for the different ages.

| Traits | Ages (weeks) |  |  |  |  |  |  |  |  |
| --- | --- | --- | --- | --- | --- | --- | --- | --- | --- |
|  | 70 | 71 | 72 | 80 | 81 | 82 | 90 | 91 | 92 |
| Genetic variances |  |  |  |  |  |  |  |  |  |
| BW | 13810 ± 2843 | 13891 ± 2849 | 13979 ± 2857 | 14891 ± 3034 | 15031 ± 3070 | 15178 ± 3108 | 16562 ± 3523 | 16762 ± 3588 | 16968 ± 3656 |
| DBWV |  | 1.59 ± 0.59 |  |  | 0.1 ± 0.14 |  |  | 1.34 ± 0.56 |  |
| LR | 16 ± 4.6 | 15.6 ± 4.4 | 15.2 ± 4.3 | 15.1 ± 4.1 | 15.4 ± 4.2 | 15.7 ± 4.4 | 21.3 ± 6.3 | 22.3 ± 6.7 | 23.3 ± 7.1 |
| MEW | 7 ± 1.6 | 7 ± 1.6 | 7.1 ± 1.6 | 7.1 ± 1.6 | 7.1 ± 1.6 | 7.1 ± 1.6 | 7.3 ± 1.7 | 7.3 ± 1.8 | 7.3 ± 1.8 |
| EGGM | 837 ± 196 | 811 ± 189 | 786 ± 183 | 614 ± 147 | 597 ± 145 | 581 ± 143 | 482 ± 146 | 473 ± 148 | 466 ± 151 |
| DFI | 47.7 ± 13 | 46.5 ± 12.5 | 45.5 ± 12.1 | 49.5 ± 12.2 | 51.4 ± 12.6 | 53.7 ± 13.2 | 83.2 ± 20.6 | 88.3 ± 21.9 | 93.7 ± 23.3 |
| FCR | 0.012 ± 0.003 | 0.012 ± 0.003 | 0.012 ± 0.003 | 0.01 ± 0.002 | 0.01 ± 0.002 | 0.01 ± 0.002 | 0.01 ± 0.003 | 0.011 ± 0.003 | 0.011 ± 0.003 |
| RFI |  | 14736 ± 3394 |  |  | 6949 ± 1794 |  |  | 13407 ± 3259 |  |
| HU | 17 ± 3.8 | 17.6 ± 3.9 | 18.4 ± 4.1 | 28.7 ± 5.8 | 30.5 ± 6.2 | 32.4 ± 6.5 | 51.2 ± 10.3 | 54.1 ± 10.9 | 57 ± 11.5 |
| YP | 1.62 ± 0.33 | 1.62 ± 0.33 | 1.61 ± 0.33 | 1.62 ± 0.32 | 1.63 ± 0.32 | 1.64 ± 0.33 | 1.79 ± 0.37 | 1.81 ± 0.37 | 1.84 ± 0.38 |
| ESS | 99963 ± 28678 | 95396 ± 27421 | 91093 ± 26256 | 66164 ± 20255 | 64235 ± 19928 | 62570 ± 19695 | 58748 ± 21250 | 59457 ± 21865 | 60430 ± 22572 |
| ESSStiff | 23038 ± 5652 | 22905 ± 5568 | 22793 ± 5496 | 22679 ± 5320 | 22762 ± 5349 | 22868 ± 5389 | 24491 ± 6109 | 24791 ± 6248 | 25114 ± 6397 |
| ESC | 25.4 ± 5.8 | 25.1 ± 5.7 | 24.8 ± 5.6 | 22.8 ± 5.2 | 22.6 ± 5.1 | 22.4 ± 5.1 | 20.8 ± 5.2 | 20.6 ± 5.2 | 20.5 ± 5.3 |
| Permanent environment variances |  |  |  |  |  |  |  |  |  |
| BW | 12256 ± 1867 | 12176 ± 1862 | 12126 ± 1860 | 12840 ± 1953 | 13069 ± 1979 | 13328 ± 2007 | 16511 ± 2339 | 17048 ± 2394 | 17616 ± 2452 |
| DBWV |  | 0.11 ± 0.96 |  |  | 0.13 ± 0.28 |  |  | 0.16 ± 0.87 |  |
| LR | 16.6 ± 3.9 | 17.1 ± 3.7 | 17.7 ± 3.6 | 22.6 ± 3.2 | 23.3 ± 3.2 | 23.9 ± 3.3 | 29.8 ± 5 | 30.6 ± 5.3 | 31.4 ± 5.6 |
| MEW | 6.3 ± 1.1 | 6.3 ± 1 | 6.3 ± 1 | 7 ± 1 | 7.2 ± 1.1 | 7.4 ± 1.1 | 9.9 ± 1.2 | 10.4 ± 1.3 | 10.8 ± 1.3 |
| EGGM | 428 ± 133 | 451 ± 128 | 474 ± 124 | 686 ± 105 | 715 ± 105 | 744 ± 105 | 1004 ± 128 | 1039 ± 133 | 1075 ± 139 |
| DFI | 73.6 ± 10 | 69.5 ± 9.5 | 66 ± 9.1 | 56.1 ± 8.3 | 57.2 ± 8.6 | 58.8 ± 8.9 | 90.6 ± 13.8 | 96.9 ± 14.7 | 103.8 ± 15.7 |
| FCR | 0.083 ± 0.002 | 0.064 ± 0.002 | 0.06 ± 0.002 | 0.072 ± 0.002 | 0.075 ± 0.002 | 0.073 ± 0.002 | 0.071 ± 0.003 | 0.074 ± 0.003 | 0.067 ± 0.003 |
| RFI |  | 4478 ± 2263 |  |  | 4087 ± 1772 |  |  | 4827 ± 3767 |  |
| HU | 13.1 ± 2.7 | 13.2 ± 2.7 | 13.3 ± 2.7 | 16 ± 3.6 | 16.5 ± 3.8 | 17.1 ± 4 | 23.5 ± 6.3 | 24.5 ± 6.7 | 25.6 ± 7 |
| YP | 1 ± 0.21 | 0.99 ± 0.21 | 0.98 ± 0.21 | 0.99 ± 0.2 | 1 ± 0.2 | 1.01 ± 0.2 | 1.17 ± 0.23 | 1.19 ± 0.24 | 1.22 ± 0.25 |
| ESS | 131350 ± 21834 | 131080 ± 20907 | 130950 ± 20063 | 134740 ± 16345 | 135820 ± 16261 | 137030 ± 16260 | 151570 ± 19272 | 154000 ± 20022 | 156550 ± 20854 |
| ESSStiff | 25123 ± 4063 | 24619 ± 3966 | 24179 ± 3881 | 22925 ± 3655 | 23052 ± 3682 | 23242 ± 3720 | 27033 ± 4456 | 27790 ± 4603 | 28611 ± 4761 |
| ESC | 26.4 ± 4.1 | 25.7 ± 4 | 25.2 ± 3.9 | 23.6 ± 3.5 | 23.7 ± 3.5 | 24 ± 3.5 | 28.8 ± 3.9 | 29.8 ± 4 | 30.8 ± 4.1 |

| Residual Variances |  |  |  |  |  |  |  |  |  |
| --- | --- | --- | --- | --- | --- | --- | --- | --- | --- |
| BW | 4763 ± 298 |  | 2835 ± 206 | 2067 ± 153 |  | 4654 ± 241 | 4968 ± 298 |  | 3572 ± 282 |
| DBWV |  | 12.48 ± 1 |  |  | 10.22 ± 0.51 |  |  | 9.51 ± 0.88 |  |
| LR | 81.8 ± 4.4 | 93.7 ± 4.8 | 86.4 ± 4.4 | 81.2 ± 4.1 | 94.9 ± 4.8 | 101.8 ± 5.1 | 96.1 ± 5.2 | 107.4 ± 5.7 | 99.3 ± 5.5 |
| MEW | 7 ± 0.4 | 5.1 ± 0.3 | 5.8 ± 0.3 | 6.7 ± 0.3 | 5.8 ± 0.3 | 6.1 ± 0.3 | 6.7 ± 0.4 | 6.4 ± 0.4 | 5.9 ± 0.4 |
| EGGM | 1868 ± 101 | 1832 ± 97 | 1593 ± 86 | 2231 ± 113 | 2428 ± 122 | 2407 ± 121 | 1748 ± 100 | 2285 ± 125 | 2381 ± 131 |
| DFI | 156.4 ± 8.5 | 90.1 ± 5.5 | 96.5 ± 5.6 | 114.6 ± 5.7 | 168.3 ± 8 | 189.4 ± 9 | 87 ± 4.7 | 40.8 ± 3.1 | 79.2 ± 4.7 |
| FCR | 0.012 ± 0.004 | 0.012 ± 0.003 | 0.012 ± 0.003 | 0.01 ± 0.004 | 0.01 ± 0.004 | 0.01 ± 0.004 | 0.01 ± 0.004 | 0.011 ± 0.004 | 0.011 ± 0.004 |
| RFI |  | 14225 ± 2095 |  |  | 24837 ± 1465 |  |  | 3751 ± 2168 |  |
| HU | 27.5 ± 1.2 | 49.8 ± 1.7 | 21.1 ± 1.2 |  |  |  | 46.6 ± 2.2 | 31.6 ± 1.5 | 47.8 ± 2.7 |
| YP | 1.34 ± 0.06 | 2.31 ± 0.08 | 1.76 ± 0.09 |  |  |  | 1.37 ± 0.07 | 1.24 ± 0.06 | 1.21 ± 0.08 |
| ESS | 308649 ± 13414 | 368380 ± 13333 | 309921 ± 16725 |  |  |  | 349216 ± 17339 | 329777 ± 15893 | 326398 ± 19866 |
| ESSstiff | 45201 ± 1965 | 45715 ± 1698 | 50575 ± 2746 |  |  |  | 52614 ± 2669 | 52915 ± 2604 | 37976 ± 2413 |
| ESC | 38.4 ± 1.6 | 29.9 ± 1.1 | 47.4 ± 2.4 |  |  |  | 37.6 ± 1.8 | 39.3 ± 1.8 | 39 ± 2.3 |
| Heritabilities |  |  |  |  |  |  |  |  |  |
| BW | 0.45 ± 0.08 |  | 0.48 ± 0.08 | 0.49 ± 0.08 |  | 0.46 ± 0.08 | 0.44 ± 0.08 |  | 0.44 ± 0.08 |
| DBWV |  | 0.12 ± 0.04 |  |  | 0.01 ± 0.01 |  |  | 0.12 ± 0.05 |  |
| LR | 0.14 ± 0.04 | 0.12 ± 0.03 | 0.13 ± 0.03 | 0.13 ± 0.03 | 0.12 ± 0.03 | 0.11 ± 0.03 | 0.14 ± 0.04 | 0.14 ± 0.04 | 0.15 ± 0.04 |
| MEW | 0.35 ± 0.07 | 0.38 ± 0.07 | 0.37 ± 0.07 | 0.34 ± 0.07 | 0.35 ± 0.07 | 0.35 ± 0.07 | 0.3 ± 0.06 | 0.3 ± 0.06 | 0.3 ± 0.07 |
| EGGM | 0.27 ± 0.06 | 0.26 ± 0.06 | 0.28 ± 0.06 | 0.17 ± 0.04 | 0.16 ± 0.04 | 0.16 ± 0.04 | 0.15 ± 0.04 | 0.12 ± 0.04 | 0.12 ± 0.04 |
| DFI | 0.17 ± 0.04 | 0.23 ± 0.06 | 0.22 ± 0.05 | 0.22 ± 0.05 | 0.19 ± 0.04 | 0.18 ± 0.04 | 0.32 ± 0.07 | 0.39 ± 0.08 | 0.34 ± 0.07 |
| FCR | 0.13 ± 0.03 | 0.16 ± 0.04 | 0.16 ± 0.04 | 0.11 ± 0.03 | 0.11 ± 0.02 | 0.11 ± 0.03 | 0.11 ± 0.03 | 0.11 ± 0.03 | 0.12 ± 0.03 |
| RFI |  | 0.27 ± 0.08 |  |  | 0.29 ± 0.04 |  |  | 0.75 ± 0.09 |  |
| HU | 0.29 ± 0.06 | 0.22 ± 0.04 | 0.35 ± 0.07 |  |  |  | 0.42 ± 0.07 | 0.49 ± 0.08 | 0.44 ± 0.07 |
| YP | 0.41 ± 0.07 | 0.33 ± 0.06 | 0.37 ± 0.06 |  |  |  | 0.41 ± 0.07 | 0.43 ± 0.07 | 0.43 ± 0.07 |
| ESS | 0.19 ± 0.05 | 0.16 ± 0.04 | 0.17 ± 0.05 |  |  |  | 0.11 ± 0.04 | 0.11 ± 0.04 | 0.11 ± 0.04 |
| ESSstiff | 0.25 ± 0.05 | 0.25 ± 0.05 | 0.23 ± 0.05 |  |  |  | 0.24 ± 0.05 | 0.24 ± 0.05 | 0.27 ± 0.06 |
| ESC | 0.28 ± 0.06 | 0.31 ± 0.06 | 0.25 ± 0.05 |  |  |  | 0.24 ± 0.05 | 0.23 ± 0.05 | 0.23 ± 0.05 |

**Supplementary Figure 1:**
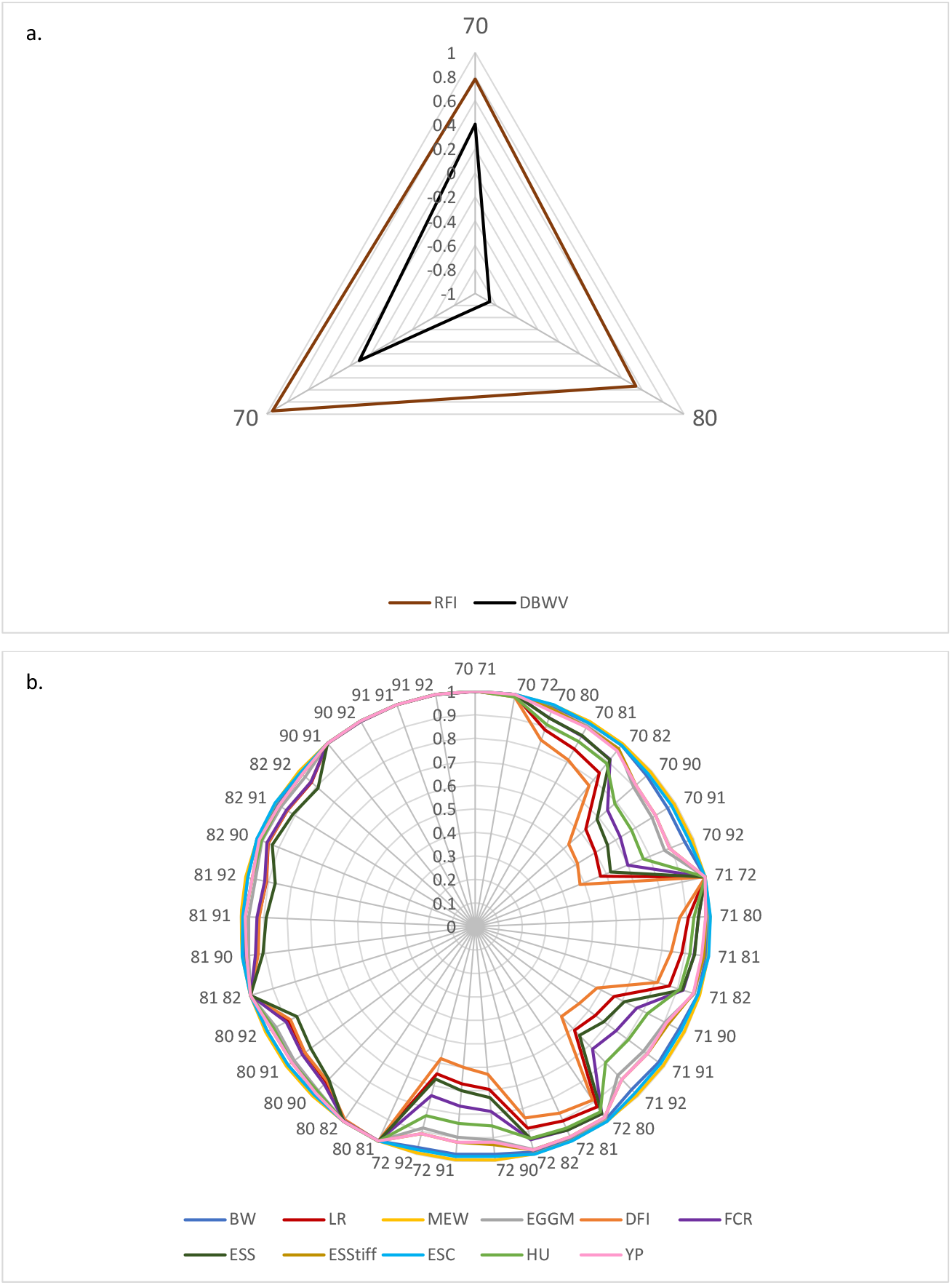
Genetic correlations between different ages of the three studied periods for DBWV and RFI (a) and the remaining traits (b).

**Supplementary Figure 2:**
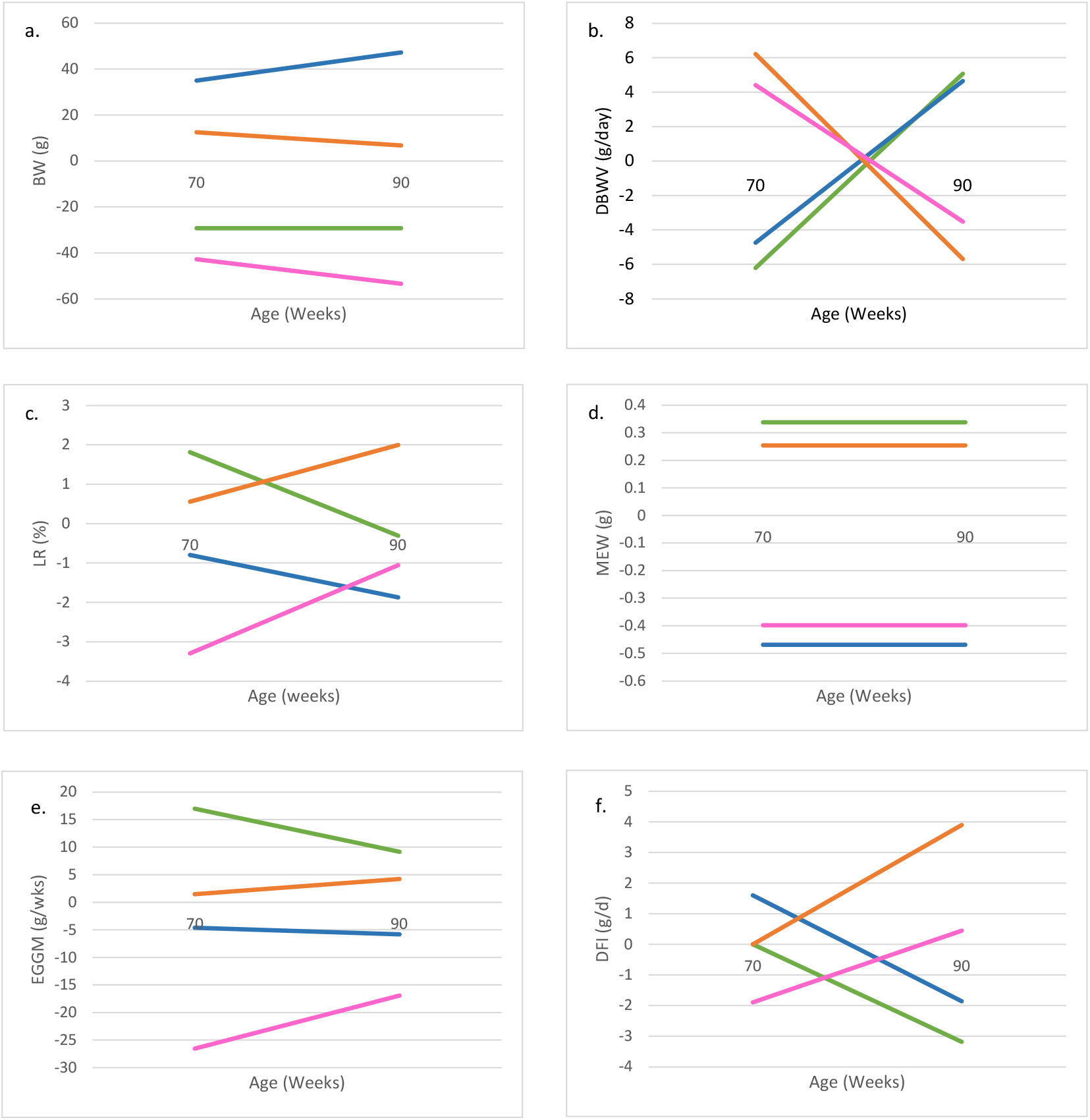
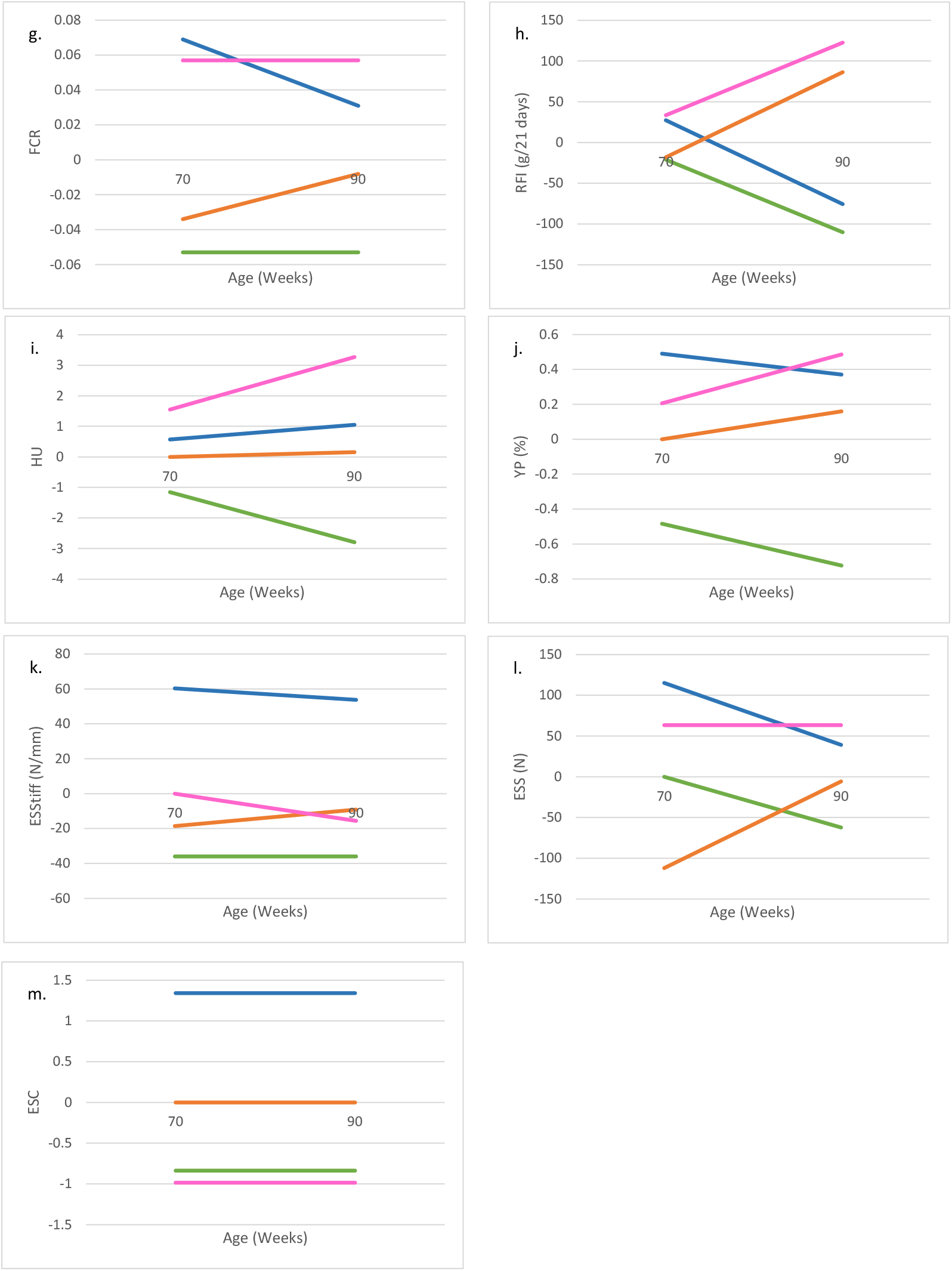
EBV evolution between 70 and 90 weeks of age for BW(a), DBWV (b), LR (c), MEW (d), EGGM (e), DFI (f), FCR(g), RFI (h), HU (i), YP (j), ESS (k), ESStiff (l), ESC (m). Green lines represent the cluster 1, blue lines the cluster2, orange line the cluster3 and pink lines the cluster4.

## References

1 OECD, Food and Agriculture Organization of the United Nations. OECD-FAO Agricultural Outlook 2022-2031. OECD; 2022.

2 Molnár A, Maertens L, Ampe B, Buyse J, Kempen I, Zoons J, et al. Changes in egg quality traits during the last phase of production: is there potential for an extended laying cycle? British Poultry Science. 2016;57:842–7.

3 Bain MM, Nys Y, Dunn IC. Increasing persistency in lay and stabilising egg quality in longer laying cycles. What are the challenges? British Poultry Science. 2016;57:330–8.

4 Livestock populations in England at 1 June 2022. GOV.UK. https://www.gov.uk/government/statistics/livestock-populations-in-england/livestock-populations-in-england-at-1-june-2022. xAccessed 21 Mar 2023.

5 Abín R, Laca A, Laca A, Díaz M. Environmental assesment of intensive egg production: A Spanish case study. Journal of Cleaner Production. 2018;179:160–8.

6 Gautron J, Réhault-Godbert S, Van de Braak TGH, Dunn IC. Review: What are the challenges facing the table egg industry in the next decades and what can be done to address them? Animal. 2021;15:100282.

7 Molnár A, Buyse J, Maertens L, Zoons J, Delezie E. Extended laying cycle of laying hens: a field study on the current situation in Belgium. 2014.

8 Rios RL, Bertechini AG, Carvalho JCC, Castro SF, Costa VA. Effect of Cage Density on the Performance of 25-to 84-Week-Old Laying Hens. Braz J Poult Sci. 2009;11:257–62.

9 Alfonso-Carrillo C, Benavides-Reyes C, de los Mozos J, Dominguez-Gasca N, Sanchez-Rodríguez E, Garcia-Ruiz AI, et al. Relationship between Bone Quality, Egg Production and Eggshell Quality in Laying Hens at the End of an Extended Production Cycle (105 Weeks). Animals. 2021;11:623.

10 Bourin M, Travel A, Souchet C, Bernard J, Ledru E, Bernardet N, et al. Effects of lengthening the laying period of laying hens on egg quality. Animal - science proceedings. 2022;13:633–4.

11 Curtis PA, Kerth LK, Anderson KE. Impact of strain on egg quality and composition during a single production cycle. Poult Sci. 2005;84:78–78.

12 Wolc A, Arango J, Settar P, O’Sullivan NP, Olori VE, White MS, et al. Genetic parameters of egg defects and egg quality in layer chickens. Poult Sci. 2012;91:1292–8.

13 Bish CL, Beane WL, Ruszler PL, Cherry JA. Body Weight Influence on Egg Production. Poultry Science. 1985;64:2259–62.

14 Parkinson GB, Roberts JR, Horn R. Pullet and layer flock uniformity, persistency and longevity: An epidemilogical, industry - based approach to improve feed efficiency: A report for the Australian Egg Corporation Limited. 2015.

15 Anderson KE, Havenstein GB, Jenkins PK, Osborne J. Changes in commercial laying stock performance, 1958-2011: thirty-seven flocks of the North Carolina random sample and subsequent layer performance and management tests. World’s Poultry Science Journal. 2013;69:489–514.

16 Venturini GC, Savegnago RP, Nunes BN, Ledur MC, Schmidt GS, El Faro L, et al. Genetic parameters and principal component analysis for egg production from White Leghorn hens. Poult Sci. 2013;92:2283–9.

17 Biscarini F, Bovenhuis H, Ellen ED, Addo S, van Arendonk JAM. Estimation of heritability and breeding values for early egg production in laying hens from pooled data. Poultry Science. 2010;89:1842–9.

18 Nurgiartiningsih VMA, Mielenz N, Preisinger R, Schmutz M, Schuler L. Heritabilities and genetic correlations for monthly egg production and egg weight of White Leghorn hens estimated based on hen-housed and survivor production. Arch Geflugelkd. 2005;69:98–102.

19 Wolc A, White IMS, Hill WG, Olori VE. Inheritance of hatchability in broiler chickens and its relationship to egg quality traits. Poultry Science. 2010;89:2334–40.

20 Wolc A, Arango J, Settar P, O’Sullivan NP, Dekkers JCM. Evaluation of egg production in layers using random regression models. Poultry Science. 2011;90:30–4.

21 Dunn IC, Bain M, Edmond A, Wilson PW, Joseph N, Solomon S, et al. Heritability and genetic correlation of measurements derived from acoustic resonance frequency analysis; a novel method of determining eggshell quality in domestic hens. Br Poult Sci. 2005;46:280–6.

22 Sambeek F van. Longer production cycles from a genetic perspective. International Poultry Production. 2011;19:27–9.

23 Grossman M, Gossman TN, Koops WJ. A model for persistency of egg production1. Poultry Science. 2000;79:1715–24.

24 Savegnago RP, Caetano SL, Ramos SB, Nascimento GB, Schmidt GS, Ledur MC, et al. Estimates of genetic parameters, and cluster and principal components analyses of breeding values related to egg production traits in a White Leghorn population. Poultry Science. 2011;90:2174–88.

25 Anang A, Mielenz N, Schüler L. Monthly model for genetic evaluation of laying hens II. Random regression. British Poultry Science. 2002;43:384–90.

26 Kirkpatrick M, Lofsvold D, Bulmer M. Analysis of the inheritance, selection and evolution of growth trajectories. Genetics. 1990;124:979–93.

27 Schaeffer LR, Jamrozik J, Kistemaker GJ, Van Doormaal J. Experience with a Test-Day Model. Journal of Dairy Science. 2000;83:1135–44.

28 Schaeffer LR, Dekkers JCM. Random regressions in animal models for test-day production in dairy cattle. Proceedings of the World Congress on Genetics applied to Livestock Production. 1994;18. Genetics and Breeding of Sheep and Goats; Breeding objectives and Breeding strategies; Genetic Parameters, Breeding Values: 443–6.

29 Jamrozik J, Schaeffer LR, Dekkers JCM. Genetic Evaluation of Dairy Cattle Using Test Day Yields and Random Regression Model. Journal of Dairy Science. 1997;80:1217–26.

30 Berry DP, Buckley F, Dillon P, Evans RD, Rath M, Veerkamp RF. Estimation of genotype×environment interactions, in a grass-based system, for milk yield, body condition score, and body weight using random regression models. Livestock Production Science. 2003;83:191–203.

31 Calus MPL, Veerkamp RF. Estimation of Environmental Sensitivity of Genetic Merit for Milk Production Traits Using a Random Regression Model. Journal of Dairy Science. 2003;86:3756–64.

32 Martin P, Ducrocq V, Gordo DGM, Friggens NC. A new method to estimate residual feed intake in dairy cattle using time series data. animal. 2021;15:100101.

33 Olori V, Hill W, Brotherstone S. The structure of the residual error variance of test day milk yield in random regression model. Workshop on Computational Breeding. 1999;20.

34 Venturini GC, Grossi DA, Ramos SB, Cruz VAR, Souza CG, Ledur MC, et al. Estimation of genetic parameters for partial egg production periods by means of random regression models. Genet Mol Res. 2012;11:1819–29.

35 Mookprom S, Boonkum W, Kunhareang S, Siripanya S, Duangjinda M. Genetic evaluation of egg production curve in Thai native chickens by random regression and spline models. Poultry Science. 2017;96:274–81.

36 Byerly TC, Kessler JW, Gous RM, Thomas OP. Feed Requirements for Egg Production1. Poultry Science. 1980;59:2500–7.

37 Bordas A, Tixier-Boichard M, Mérat P. Direct and correlated responses to divergent selection for residual food intake in Rhode Island Red laying hens. Br Poult Sci. 1992;33:741–54.

38 R Core Team. R: A language and environment for statistical computing. Vienna, Austria; 2020.

39 Haugh H. The Haugh Unit for Measuring Egg Quality. The US Egg & Poultry Magazine. 1937;43:552–5.

40 Falconer DS, Mackay TFC. Introduction to quantitative genetics. 4th ed. Harlow, England: Prentice Hall; 1996.

41 Finlay KW, Wilkinson GN. The analysis of adaptation in a plant-breeding programme. Aust J Agric Res. 1963;14:742–54.

42 Gilmour AR, Gogel BJ, Cullis BR, Welham SJ, Thompson R. ASReml User Guide Release 4.2 Functional Specification. VSN International Ltd, VSN International Ltd. Hemel Hempstead. UK; 2021.

43 Altman DG, Bland JM. How to obtain the P value from a confidence interval. BMJ. 2011;343:d2304.

44 Lê S, Josse J, Husson F. FactoMineR: An R Package for Multivariate Analysis. J Stat Soft. 2008;25:1–18.

45 Molnár A, Maertens L, Ampe B, Buyse J, Zoons J, Delezie E. Effect of different split-feeding treatments on performance, egg quality, and bone quality of individually housed aged laying hens. Poultry Science. 2018;97:88–101.

46 Anang A, Mielenz N, Schüler L. Genetic and phenotypic parameters for monthly egg production in White Leghorn hens. Journal of Animal Breeding and Genetics. 2008;117:407–15.

47 Luo PT, Yang RQ, Yang N. Estimation of Genetic Parameters for Cumulative Egg Numbers in a Broiler Dam Line by Using a Random Regression Model. Poultry Science. 2007;86:30–6.

48 Wolc A, Szwaczkowski T. Estimation of genetic parameters for monthly egg production in laying hens based on random regression models. J Appl Genetics. 2009;50:41–6.

49 Freitas L, Tinôco I, Baêta F, Barbari M, Conti L, Júnior C, et al. Correlation between egg quality parameters, housing thermal conditions and age of laying hens. Agronomy Research. 2007;15:687–93.

50 Silversides FG, Budgell K. The Relationships Among Measures of Egg Albumen Height, pH, and Whipping Volume. Poultry Science. 2004;83:1619–23.

51 Marzec A, Damaziak K, Kowalska H, Riedel J, Michalczuk M, Koczywąs E, et al. Effect of Hens Age and Storage Time on Functional and Physiochemical Properties of Eggs. Journal of Applied Poultry Research. 2019;28:290–300.

52 Bar A, Razaphkovsky V, Vax E. Re-evaluation of calcium and phosphorus requirements in aged laying hens. British Poultry Science. 2002;43:261–9.

53 Rodriguez-Navarro A, Kalin O, Nys Y, Garcia-Ruiz JM. Influence of the microstructure on the shell strength of eggs laid by hens of different ages. Br Poult Sci. 2002;43:395–403.

54 Dunn IC, De Koning D-J, McCormack HA, Fleming RH, Wilson PW, Andersson B, et al. No evidence that selection for egg production persistency causes loss of bone quality in laying hens. Genet Sel Evol. 2021;53:11.

55 Wolc A, Arango J, Settar P, Fulton JE, O’Sullivan NP, Preisinger R, et al. Genome-wide association analysis and genetic architecture of egg weight and egg uniformity in layer chickens. Anim Genet. 2012;43:87–96.

56 Koch RM, Swiger LA, Chambers D, Gregory KE. Efficiency of Feed Use in Beef Cattle. J Anim Sci. 1963;22:486–94.

57 Kumar D, Raginski C, Schwean-Lardner K, Classen HL. Assessing the response of hen weight, body composition, feather score, egg quality, and level of excreta nitrogen content to digestible balanced protein intake of laying hens. Can J Anim Sci. 2018;98:619–30.

58 Dunn I. 11 - Poultry breeding for egg quality: traditional and modern genetic approaches. In: Nys Y, Bain M, Van Immerseel F, editors. Improving the Safety and Quality of Eggs and Egg Products. Woodhead Publishing; 2011. p. 245–60.

59 Zhang L-C, Ning Z-H, Xu G-Y, Hou Z-C, Yang N. Heritabilities and genetic and phenotypic correlations of egg quality traits in brown-egg dwarf layers. Poultry Science. 2005;84:1209–13.

60 Icken W, Looft C, Schellander K, Cavero D, Blanco A, Schmutz M, et al. Implications of genetic selection on yolk proportion on the dry matter content of eggs in a White Leghorn population. Br Poult Sci. 2014;55:291–7.

61 Liu Z, Sun C, Yan Y, Li G, Shi F, Wu G, et al. Genetic variations for egg quality of chickens at late laying period revealed by genome-wide association study. Sci Rep. 2018;8:10832.

62 Bécot L, Bédère N, Ferry A, Burlot T, Le Roy P. Egg production in nests and nesting behaviour: genetic correlations with egg quality and BW for laying hens on the floor. animal. 2023;17:100958.

63 Yuan J, Dou T, Ma M, Yi G, Chen S, Qu L, et al. Genetic parameters of feed efficiency traits in laying period of chickens. Poultry Science. 2015;94:1470–5.

64 Wolc A, Arango J, Jankowski T, Settar P, Fulton JE, O’Sullivan NP, et al. Pedigree and genomic analyses of feed consumption and residual feed intake in laying hens. Poultry Science. 2013;92:2270–5.

65 Tixier-Boichard M, Boichard D, Groeneveld E, Bordas A. Restricted maximum likelihood estimates of genetic parameters of adult male and female Rhode Island red chickens divergently selected for residual feed consumption. Poult Sci. 1995;74:1245–52.

66 Zhou Q, Lan F, Gu S, Li G, Wu G, Yan Y, et al. Genetic and microbiome analysis of feed efficiency in laying hens. Poult Sci. 2022;102:102393.

67 Aggrey SE, Karnuah AB, Sebastian B, Anthony NB. Genetic properties of feed efficiency parameters in meat-type chickens. Genet Sel Evol. 2010;42:25–30.

68 Kavlak AT, Uimari P. Inheritance of feed intake-based resilience traits and their correlation with production traits in Finnish pig breeds. J Anim Sci. 2024;102.

69 Casto-Rebollo C, Nuñez P, Gol S, Reixach J, Ibáñez-Escriche N. Variability of daily feed intake as an indicator of resilience in Pietrain pigs. animal. 2025;19:101415.

70 Ni A, Calus MPL, Bovenhuis H, Yuan J, Wang Y, Sun Y, et al. Genetic parameters, reciprocal cross differences, and age-related heterosis of egg-laying performance in chickens. Genetics Selection Evolution. 2023;55:87.

71 Alipanah M, Deljo J, Rokouie M, Mohammadnia R. HERITABILITIES AND GENETIC AND PHENOTYPIC CORRELATIONS OF EGG QUALITY TRAITS IN KHAZAK LAYERS. Trakia Journal of Sciences. 2013;11.

72 Wolc A, Drobik-Czwarno W, Jankowski T, Arango J, Settar P, Fulton JE, et al. Accuracy of genomic prediction of shell quality in a White Leghorn line. Poult Sci. 2020;99:2833–40.

73 Romé H, Varenne A, Hérault F, Chapuis H, Alleno C, Dehais P, et al. GWAS analyses reveal QTL in egg layers that differ in response to diet differences. Genetics Selection Evolution. 2015;47:83.

74 Knaga S, Kibała L, Kasperek K, Rozempolska-Rucińska I, Buza M, Zięba G. Eggshell strength in laying hens’ breeding goals - a review. Animal Science Papers & Reports. 2019;37:119–36.

75 Park J-A, Sohn S-H. The Influence of Hen Aging on Eggshell Ultrastructure and Shell Mineral Components. Korean J Food Sci Anim Resour. 2018;38:1080–91.

76 Williams KC. Some factors affecting albumen quality with particular reference to Haugh unit score. World’s Poultry Science Journal. 1992. 10.1079/WPS19920002.

77 Padhi MK, Chatterjee RN, Haunshi S, Rajkumar U. Effect of age on egg quality in chicken. Indian Journal of Poultry Science. 2013;48:122–5.

78 Schnyder U, Hofer A, Labroue F, Künzi N. Genetic parameters of a random regression model for daily feed intake of performance tested French Landrace and Large White growing pigs. Genet Sel Evol. 2001;33:635–58.

79 Miranda JC, León JM, Pieramati C, Gómez MM, Valdés J, Barba C. Estimation of Genetic Parameters for Peak Yield, Yield and Persistency Traits in Murciano-Granadina Goats Using Multi-Traits Models. Animals. 2019;9:411.

80 Podisi BK, Knott SA, Burt DW, Hocking PM. Comparative analysis of quantitative trait loci for body weight, growth rate and growth curve parameters from 3 to 72 weeks of age in female chickens of a broiler–layer cross. BMC Genet. 2013;14:22.

81 Howard JT, Jiao S, Tiezzi F, Huang Y, Gray KA, Maltecca C. Genome-wide association study on legendre random regression coefficients for the growth and feed intake trajectory on Duroc Boars. BMC Genet. 2015;16:1–11.

82 van Eck LM, Enting, H., Carvalhido, I. J., Chen, H., and Kwakkel RP. Lipid metabolism and body composition in long-term producing hens. World’s Poultry Science Journal. 2023;79:243–64.

83 Pérez-Bonilla A, Jabbour C, Frikha M, Mirzaie S, Garcia J, Mateos GG. Effect of crude protein and fat content of diet on productive performance and egg quality traits of brown egg-laying hens with different initial body weight. Poult Sci. 2012;91:1400–5.

84 van Eck LM, Enting H, Carvalhido IJ, Chen H, Kwakkel RP. Lipid metabolism and body composition in long-term producing hens. World’s Poultry Science Journal. 2023;79:243–64.

85 Summers JD, Leeson S. Laying hen performance as influenced by protein intake to sixteen weeks of age and body weight at point of lay. Poult Sci. 1994;73:495–501.

86 Ghasempour A, Ahmadi E. Assessment of environment impacts of egg production chain using life cycle assessment. Journal of Environmental Management. 2016;183:980–7.

87 Sharma MK, McDaniel CD, Kiess AS, Loar RE, Adhikari P. Effect of housing environment and hen strain on egg production and egg quality as well as cloacal and eggshell microbiology in laying hens. Poultry Science. 2022;101:101595.

